# Bipolar-associated miR-499-5p controls neuroplasticity by downregulating the Ca_v_1.2 L-type voltage gated calcium channel subunit CACNB2

**DOI:** 10.1101/2021.06.09.447782

**Authors:** Martins H.C., Sungur A.Ö., Gilardi C., Pelzl M., Bicker S., Gross F., Winterer J., Kisko T.M., Malikowska-Racia N., Braun M.D., Brosch K., Nenadic I., Stein F., Meinert S., Schwarting R.K.W., Dannlowski U., Kircher T., Wöhr M., Schratt G.

**Affiliations:** Lab of Systems Neuroscience, Institute for Neuroscience, Department of Health Science and Technology, Swiss Federal Institute of Technology ETH, 8057 Zurich, Switzerland; Institute for Physiological Chemistry, Biochemical-Pharmacological Center Marburg, Philipps-University of Marburg, 35032 Marburg, Germany; Behavioural Neuroscience, Experimental and Biological Psychology, Faculty of Psychology, Philipps-University of Marburg, 35032 Marburg, Germany; Center for Mind, Brain, and Behavior, Philipps-University of Marburg, 35032 Marburg, Germany; Social and Affective Neuroscience Research Group, Laboratory of Biological Psychology, Research Unit Brain and Cognition, Faculty of Psychology and Educational Sciences, KU Leuven, 3000 Leuven, Belgium; Leuven Brain Institute, KU Leuven, 3000 Leuven, Belgium; Dept. of Psychiatry and Psychotherapy, University of Marburg, 35039 Marburg, Germany; Institute for Translational Psychiatry, University of Münster, 48149 Münster, Germany; current address: Psychiatry and Psychotherapy, University of Tübingen, 72076 Tübingen, Germany; current address: Department of Behavioral Neuroscience and Drug Development, Maj Institute of Pharmacology, Polish Academy of Sciences, 31343 Krakow, Poland

**Keywords:** microRNA, miR-499, bipolar disorder, neuroplasticity, dendritogenesis, calcium channel, maltreatment, early life adversity, cognitive function

## Abstract

Bipolar disorder (BD) is a chronic mood disorder characterized by alternating manic and depressive episodes, often in conjunction with cognitive deficits. Dysregulation of neuroplasticity and calcium homeostasis as a result of complex genetic environment interactions are frequently observed in BD patients, but the underlying molecular mechanisms are largely unknown. Here, we show that a BD-associated microRNA, miR-499-5p, regulates neuronal dendrite development and cognitive function by downregulating the BD risk gene *CACNB2*. miR-499-5p expression is increased in peripheral blood of BD patients and healthy subjects at risk of developing the disorder due to a history of childhood maltreatment. This up-regulation is paralleled in the hippocampus of rats which underwent juvenile social isolation. Elevating miR-499-5p levels in rat hippocampal pyramidal neurons impairs dendritogenesis and reduces surface expression and activity of the voltage-gated L-type calcium channel Ca_v_1.2. We further identified *CACNB2*, which encodes a regulatory β-subunit of Ca_v_1.2, as a direct target of miR-499-5p in neurons. *CACNB2* downregulation is required for the miR-499-5p dependent impairment of dendritogenesis, suggesting that CACNB2 is an important downstream target of miR-499-5p in the regulation of neuroplasticity. Finally, elevating miR-499-5p in the hippocampus *in vivo* is sufficient to induce short-term memory impairments in rats haploinsufficient for the Ca_v_1.2 pore forming subunit *Cacna1c*. Taken together, we propose that stress-induced upregulation of miR-499-5p contributes to dendritic impairments and deregulated calcium homeostasis in BD, with specific implications for the neurocognitive dysfunction frequently observed in BD patients.

## INTRODUCTION

Bipolar disorder (BD) is a severe and chronic mood disorder defined by recurring (hypo)manic and depressive episodes. It represents a highly debilitating condition leading to cognitive impairments and an especially high risk for suicide deaths (1, 2). BD has one of the highest heritability rates among mental illnesses (3). As a result, genetic studies found important susceptibility genes but also revealed a complex and heterogeneous genetic architecture where no single gene variation by itself is sufficient to cause the disorder. Therefore, according to the current consensus, an interaction between genetic and environmental (GxE) risk factors is required for a full manifestation of the disease in affected individuals.

With regard to the genetic component, recent GWAS and analysis of *de novo* mutations found strong associations between single nucleotide polymorphisms (SNPs) in L-type voltage-gated calcium channel (LVGCC) coding genes and BD. This includes genetic variants in the CACNA1C (4) and CACNB2 (5) loci, which encode the α_1_ pore subunit and the auxiliary β subunit of the LVGCC Ca_v_1.2, respectively. Ca_v_1.2 channels are the primary mediators of depolarization-induced calcium entry into neurons. As such they play critical roles in the regulation of neuronal excitability (6), synaptic plasticity (7), learning and memory (8), and gene transcription (9). Particularly well studied is their function in the promotion of dendrite growth and arborization in response to neuronal activity (10). Thus, the convergence of several genetic associations into one common pathway implicates Ca_v_1.2 channel dysfunction as a main genetic risk factor of BD. Concerning environmental factors, early life adversity, particularly childhood abuse and neglect, is known to significantly increase the risk to develop BD and to correlate with poorer disease outcomes (11). A common cellular endpoint of GxE risk factors in BD is defective neuroplasticity, in particular reductions of dendritic arborization and synapse density in several brain areas of BD patients, including the hippocampus (12). However, the molecular pathways which integrate GxE risk factors to induce impairments in neuroplasticity during BD are largely unknown.

MicroRNAs (miRNAs) are a large family of small, non-coding RNAs which act as post-transcriptional repressors of gene expression by binding to partially complementary sequences in the 3’ untranslated region (UTR) of target mRNAs (13). In animals, miRNAs are widely expressed in the brain where they regulate various aspects of neuroplasticity, e.g. dendritogenesis and dendritic spine development (14), in an activity-dependent manner. A putative involvement of miRNAs in BD etiology is supported by several recent observations. First, differential expression of miRNAs is found in *post mortem* brain and blood of BD patients (15). Second, both circulating and brain miRNA levels are modulated by the intake of antidepressants (16) and mood stabilizers (17, 18). Third, *in vivo* manipulation of specific miRNA candidates in rodents led to phenotypes in affective-like and cognitive behaviors via the modulation of serotonin, glucocorticoid, neurotrophic factor, and Wnt signaling pathways (19). Lastly, variations in miRNA genes that confer susceptibility to BD have recently been identified (20). However, it is still unknown how the dysregulation of specific miRNAs and associated pathways impairs neuronal function and contributes to BD.

In this study, we focused on miR-499-5p, a miRNA which after correction for multiple testing showed a nominally significant association with BD in a recent GWAS (20). Interestingly, miR-499-5p plays important roles in the physiology and pathology of the cardiovascular system (21), but its function in the nervous system is completely unexplored. We found that miR-499-5p levels are strongly up-regulated in the blood of BD patients and healthy subjects at risk of developing the disorder due to a history of childhood maltreatment, as well as in the hippocampus of socially isolated rats. In hippocampal neurons, miR-499-5p targets the recently identified BD risk gene *Cacnb2* and controls dendritic development, Ca_v_1.2 surface expression, and current density. In the *Cacna1c*^+/−^ rat model, overexpression of miR-499-5p in the hippocampus induced deficits in short-term recognition memory, providing a potential link between GxE risk factors. Together, this suggests a novel mechanism whereby early life adversity induces excessive miR-499-5p expression, which in turn negatively impacts neuronal calcium homeostasis, neuroplasticity, and cognitive function.

## MATERIALS AND METHODS

### Human study

BD patients (n=26) and healthy controls (n=26), and healthy subjects with (n=17) or without (n=18) a history of childhood maltreatment were obtained from the Departments of Psychiatry at the University of Marburg and the University of Münster, Germany, as part of the FOR2107 cohort (22). The diagnosis was ascertained using the German version of the Structured Clinical Interview for DSM-IV (SCID-I) (23)and subsequently translated to ICD-10 diagnoses (24). Subjects were excluded if they had substance-related disorders, severe neurological or other medical disorders. Healthy control subjects underwent the same diagnosis procedure as the patients. Healthy subjects were included in the childhood maltreatment study if at least one subscale of the Childhood Trauma Questionnaire (CTQ) reached the threshold for maltreatment (Emotional Abuse >= 10, Physical Abuse >= 8, Sexual Abuse >= 8, Emotional Neglect >= 15, and Physical Neglect >= 8) (25). The detailed demographic and clinical data are summarized in Supplementary Tables 1 and 2. The procedures involving humans were approved by the ethics committees of the Medical Faculties of the Universities of Münster (2014-422-b-S) and Marburg (AZ: 07/14). Written informed consent was obtained from all participants before examination.

#### Human PBMC sample processing

PBMCs were isolated from 10mL of whole blood using the LeukoLOCK technology (Thermo Scientific) by the Biomaterialbank Marburg, Germany. Total RNA extraction from PBMCs was performed using the TRIzol™ Reagent (Thermo Fisher), according to the manufacturer’s protocol.

### Animal Study

#### Ethics approval

All procedures were conducted in strict accordance with the National Institutes of Health Guidelines for the Care and Use of Laboratory Animals and the relevant local or national rules and regulations of Germany and were subject to prior authorization by the local government (MR 20/35 Nr. 19/ 2014; Tierschutzbehörde, Regierungspräsdium Gieβen, Germany).

#### Cacna1c^+/−^ animal breeding

Constitutive heterozygous *Cacna1c*^+/–^ rats were generated by SAGE Labs (now Horizon Discovery Ltd.) on a Sprague Dawley background via zincfinger nucleases following a previously established protocol (26). A heterozygous breeding protocol was used to obtain offspring from both *Cacna1c*^+/−^ and *Cacna1c*^+/+^ (WT) genotypes as previously established (27). Rats were housed under standard laboratory conditions (22 ± 2°C and 40–70% humidity) with free access to standard rodent chow and water. Genotyping was performed as previously described (27).

#### Juvenile social isolation paradigm

After weaning on PND 21, male rats were socially housed in groups of 4-6 with same-sex littermate partners (Group-housed, n=9) or housed alone (Isolated, n=9), applying a previously established protocol (28). The hippocampus of the right hemispheres was removed immediately following the 4 weeks of exposure to the experimental housing conditions at ~2 months of age.

#### Rat primary cultures

Primary rat hippocampal and cortical neuronal cultures were prepared from E18 Sprague Dawley rats (Janvier Laboratories) as previously described (29). For the preparation of primary cultures from *Cacna1c*^+/−^ or WT rat embryos, the hippocampus from each embryo was dissected and collected separately for mechanical dissociation and plating while the cortex of each embryo was collected separately for genotyping. DNA was extracted from cortical tissue using the TriFast™ reagent protocol (VWR) and the *Cacna1c* gene was amplified with the PfuPlus! DNA Polymerase (Roboklon), according to the manufacturer’s instructions, using the following primers: *Cacna1c* (369 bp) forward 5’-GCTGCTGAGCCTTTTATTGG-3’ and reverse 5’-GTCAGCAGCTATCCAGGAGG-3’.

#### Drug treatments

Neuronal-enriched cultures were obtained by treating cells with fluorodeoxyuridine (FUDR; Sigma) + uridine (Sigma) (final concentration 10μM) from the *day in vitro* 3 (*DIV* 3) (glial-cell depleted). Treatment with Dexamethasone (DEX; Sigma) from DIV 5-10 to a final concentration of 500μM and re-applied every 2 days was used to activate glucocorticoid receptors (GRs) (30). To control for vehicle effects, cells were treated with DMSO to a final volume concentration of 0.1% (v/v).

#### DNA constructs

The *Cacnb2* 3’UTR (Ensemble ID: ENSG00000165995) was amplified from a rat brain cDNA library and cloned into the pmirGLO dual-luciferase expression vector (Promega) using the PfuPlus! DNA Polymerase (Roboklon). Mutation of the miR-499-5p binding site was achieved using the Pfu Plus! DNA Polymerase Site-Directed Mutagenesis Protocol (Roboklon), according to manufacturer’s instructions. For AAV-mediated overexpression of miR-499-5p *in vivo*, a chimeric miR-499 hairpin (AAV-miR-499) was generated as previously described (31) Viral vectors were produced by the Viral Vector Facility of the Neuroscience Center Zurich (https://www.vvf.uzh.ch). The vector pβA-*CACNA1C*-HA (Ca_v_1.2-HA) was kindly gifted by Prof. Amy Lee (Department of Neurology, University of Iowa, Iowa, USA). Primer sequences are provided in Supplementary Table 3.

#### Transfections

Primary hippocampal neurons were transfected with plasmid DNA, miRNA mimics (Ambion™ Pre-miR miRNA Precursor, Thermo Fisher), and pLNAs (miRCURY LNA miRNA Power Inhibitors; QIAGEN) using the Lipofectamine 2000 reagent (Thermo Fisher). To measure *Cacnb2* mRNA levels upon miR-499-5p overexpression, hippocampal neurons were transfected at DIV7 with 10nM of the miR-499-5p or control mimic (Ambion™ Pre-miR miRNA Precursor: miR-499-5p and Neg Control 1, Thermo Fisher) with the Lipofectamine RNAiMAX reagent (Thermo Fisher) and processed 7 days later for total RNA extraction. To validate the overexpression efficiency of the pMT2-*CACNB2* plasmid (Addgene, #107424), freshly isolated cortical neurons were transfected using the P3 Primary Cell 4D-Nucleofector Kit (Lonza, LZ-V4XP-3024) and the program DC-104 of the 4D-Nucleofector device (Lonza). After 5 DIV, cells were processed for Western Blot analysis.

#### Stereotaxic Brain Injections

Overexpression of miR-499-4p *in vivo* was carried out by injecting the chimeric miR-499 hairpin in the rat hippocampus as previously described (32). Male and female WT and Cacna1c+/− rats (Sprague Dawley background) with 1.5-2 months were briefly anesthetized with isoflurane and placed in a stereotaxic frame. Stereotaxic surgery was performed under isoflurane anesthesia (Baxter Deutschland GmbH). For analgesia, animals received buprenorphine (0.05 mg/kg) 30 min before surgery and a subcutaneous injection of 0.4mL local anesthetic (Xylocaine 2% with Adrenaline 1:100,000) at the site of the incision immediately before surgery. Microinjections were delivered using a 30-gauge stainless steel infusion cannula connected to a 10μL Hamilton syringe by polyethylene tube. 1μL of virus was injected in the dorsal hippocampus and 1μl of virus in the ventral hippocampus (either AAV-miR-499 or AAV-Control) over 3 min via a microinjection pump (World Precision Instruments). The infusion cannula was left in place for an additional 3 min thereafter to allow diffusion. Hippocampal injections were performed bilaterally using the following coordinates, with bregma serving as reference: Dorsal hippocampus: anteroposterior = −3mm, mediolateral = ±2mm, and dorsoventral= +3.5 mm; Ventral hippocampus: anteroposterior = −4.8mm, mediolateral = ±4.8mm, and dorsoventral= +6.4mm (33). Necessary pain management (buprenorphine) was applied and the health of the animals was evaluated post-operative over the course of 7 consecutive days. The left hippocampus was freshly snap-frozen for biochemistry and the right hemisphere was fixed in 4% paraformaldehyde (PFA)-saline followed by dehydration in 30% sucrose-PBS solution for at least 2 days each. Brains mounted in tissue mounting fluid (Tissue-Tek O.C.T Compound, Sakura Finetek Europe B.V.) were sectioned (80 μm) using a cryostat (Histocom AG) and kept in a cryoprotectant solution for long-term storage. Images were acquired in a widefield microscope (Axio ObserverZ1/7, Zeiss).

#### Single-molecule fluorescence in situ hybridization (smFISH)

smFISH for miRNA detection on hippocampal cultures was performed using the QuantiGene ViewRNA miRNA Cell Assay Kit (Thermo Fisher) according to the manufacturer’s protocol with slight modifications. To preserve dendrite morphology, protease treatment was reduced to a dilution of 1:10,000 in PBS for 45 sec. The probes hsa-miR-499a-5p (Alexa Fluor 546; Thermo Fisher) and *CamK2* (Alexa Fluor 488; Thermo Fisher) were used for smFISH.

#### Immunostaining of primary cultures

Hippocampal neurons were co-transfected with 300ng of Ca_v_1.2-HA at DIV6 and 10nM of the miR-499-5p or control mimic (AmbionTM Pre-miR miRNA Precursor: miR-499-5p and Neg Control 1, Thermo Fisher), and processed for immunostaining 13 days after. Cells were fixed in 4% PFA-4% sucrose in PBS for 15 min, rinsed with PBS, and permeabilized in 10% normal goat serum (NGS; vol/vol) containing 0.1% Triton (vol/vol) for 10 min. Blocking was performed for 30 min in 10% NGS. Cells were then sequentially labeled with Anti-HA High Affinity (1:100, Roche) and Anti-Rat, Alexa Fluor 546 (1:4000, Thermo Fisher), both diluted in 10% NGS. After three washes with PBS, coverslips were mounted on microscope slides using AquaPoly/mount (Polysciences Inc.). Surface staining of Ca_v_1.2-HA clusters was performed as previously described (34). Images were acquired in a confocal laser-scanning microscope (CLSM 880, Zeiss) and analyzed with a custom Python-script in the context of the ImageJ framework freely available via the ImageJ-update site (https://github.com/dcolam/Cluster-Analysis-Plugin).

#### Sholl Analysis

Hippocampal neurons were transfected at DIV 5 with 100ng of GFP alone or co-transfected with 10nM of the miR-499-5p or control mimic (AmbionTM Pre-miR miRNA Precursor: miR-499-5p and Neg Control 1, Thermo Fisher). After 5 days of expression, the cells were fixed using 4% PFA-4% sucrose in PBS for 15 min, washed three times in PBS, and mounted on coverslips using AquaPoly/Mount (Chemie Brunschwig). Images were acquired in a widefield microscope (Axio ObserverZ1/7, Zeiss) and Sholl analysis was performed using the “Sholl Analysis” ImageJ plugin (35).

#### RT-qPCR

Total RNA extraction from brain tissue and primary cultures was performed using the TRIzol™ Reagent (Thermo Fisher), per manufacturer’s instructions. RNA samples were first treated with the TURBO DNase enzyme (Thermo Fisher). To detect mRNAs, total RNA was reverse transcribed with the iScript cDNA synthesis kit (Bio-Rad) and RT-qPCR was performed using the iTaq SYBR Green Supermix with ROX (Bio-Rad) on the CFX384 Real-Time System (BioRad). To detect miRNAs, the TaqMan MicroRNA Reverse Transcription Kit (Thermo Fisher) and the TaqMan Universal PCR Master Mix (Thermo Fisher) were used, according to the manufacturer’s instructions. Data were analyzed according to the ΔΔCt method, normalized first to the U6 snRNA housekeeping gene. Primer sequences are provided in Supplementary Table 3.

#### Western Blot

Proteins from hippocampal tissue were isolated from the phenol-ethanol supernatant saved from the RNA isolation using the TRIzol™ Reagent (Thermo Fisher) protocol, according to the manufacturer’s directions. From neuronal cultures, protein extracts were obtained by lysing cells in lysis buffer (50 mM Tris-pH: 7.5, 150 mM NaCl, 1% Triton-X100, 1x Complete Protease Inhibitor Cocktail (Roche)). Typically, 10μg of protein sample mixed with 4x Laemmli Sample Buffer (Bio-Rad) were separated on a 10% SDS-PAGE. After electrophoresis, proteins were transferred to a nitrocellulose membrane using a Trans-Blot Turbo system (Bio-Rad), according to the manufacturer’s protocol. Membranes were blocked in 5% milk prepared in Tris-buffered saline containing 0.1% Tween20 and incubated in primary antibody solution overnight. Antibody dilutions (rabbit anti-*CACNB2* (1:1000, Abcam), rabbit anti-*GAPDH* (1:2000, Millipore), and mouse anti-*GFP* (1:1000, Novus Biological)) were prepared in blocking solution. Membranes were washed 5 times in 5% milk before incubation in HRP (horseradish peroxidase)-conjugated secondary antibody prepared in blocking solution for 1h. Following incubation, membranes were washed 5 times in TBS-T, developed with the Clarity™ Western ECL Substrate (Bio-Rad) according to manufacturer’s guidelines, and visualized with the ChemiDocTM MP, Imaging System (BioRad).

#### Luciferase Reporter Assay

Hippocampal neurons were co-transfected at DIV 5 with 10nM of the miR-499-5p or control mimic (AmbionTM Pre-miR miRNA Precursor: miR-499-5p or Neg Control 1, Thermo Fisher) and 100ng of the *Cacnb2* 3’UTR luciferase reporters. To evaluate the effect of the miR-499-5p knockdown, neurons were co-transfected at DIV 19 with 10nM of a miR-499-5p or control pLNA inhibitor (miRCURY LNA miRNA-499-5p Power Inhibitor: miR-499-5p or Neg Control A, QIAGEN). After 72h of expression, cell lysis and luciferase assay were performed using the Dual-Luciferase Reporter Assay System (Promega), following a modified protocol (36). Luciferase activity was measured on the GloMax Discover GM3000 (Promega), according to the manufacturer’s instructions.

#### Electrophysiology

For the recording of ICa,L from primary cultured hippocampal neurons (DIV 15-16), transfected with the miR-499-5p or control mimic (AmbionTM Pre-miR miRNA Precursor: miR-499-5p or Neg Control 1, Thermo Fisher) for 5-6 days, whole-cell patch-clamp was performed as previously described (37). The extracellular solution was composed of (in mM) 110 NaCl, 2.5 KCl, 15 HEPES, 2 CaCl_2_, 2 MgCl_2_, 20 glucose, 20 TEA-Cl and 5 4-AP (adjusted to pH 7.3 with NaOH) and intracellular solution 125 CsCl, 20 TEA-Cl, 0.5 EGTA, 10 HEPES, 4 Mg-ATP, 0.3 GTP (adjusted to pH 7.3 with CsOH). 1μM TTX, 1μM Gabazine, 10μM CNQX and 1μM ω-conotoxin (CTx) MVIIC, 1μM ω-CTxGVIA, and 30μM niflumic acid were added to the extracellular buffer to block to synaptic transmission, P/Q-type and N-type Ca^2+^ channels and Ca^2+^-activated Cl^−^ channels, respectively. Cells were held at −60 mV to inactivate T-type Ca^2+^ channels. Leak and capacitive currents were subtracted using the P/4 method. Series resistance compensation was enabled in all experiments (compensation 50–70%). The sampling frequency was 50 kHz and the filter frequency 10kHz. For Ca^2+^ activation curves, Ca^2+^ currents were obtained by depolarizing cells from −60 mV to potentials between −60 and +60 mV (in 5 mV increments). For Ca^2+^ inactivation, cells were held from −70mV to 0mV (in 5mV increments), and after a depolarizing step to 10mV cells were hyperpolarized to −70mV. Peak Ca2+ currents were normalized to membrane capacitance and were plotted as a function of the membrane potential.

#### Novel object recognition test

Recognition memory was assessed in male and female WT and *Cacna1c*^+/−^ rats approximately one week after injection of the chimeric miR-499-5p hairpin. Behavioral experiments were carried out during the light phase of a 12:12 h light/dark cycle (lights on at 07:00 h). Rats were handled for three consecutive days before behavioral testing in a standardized way for 5 min. The novel object recognition test was performed as previously described (38). Behavioral analysis was performed by an experienced observer blind to experimental conditions. In total 7 WT animals and 3 *Cacna1c*^+/−^ animals were excluded from final statistical analyses since miR-499-5p expression did not exceed the mean expression of miR-499-5p + 2x s.d. of control animals (Supplementary Figure 1 and 2).

#### Statistical Analysis

All statistical tests were performed using GraphPad Prism version 8.2.0 for Windows (GraphPad Software, San Diego, California USA, www.graphpad.com). The number of independent experiments is indicated in the plots. Bar graphs represent mean ± s.d. unless stated otherwise. Boxplots represent median (box: two quantiles around the median; whiskers: Tukey; point outside: Outliers). Normality was tested using the Shapiro-Wilk test. Normally distributed data were tested using one or two sample Student’s T-test (always two-sided) or ANOVA followed by post hoc Tukey test and otherwise for nonnormal data the non-parametric test Mann-Whitney-U test. Correlations were calculated using the Spearman correlation coefficient with two-tailed analysis. Significant changes in the BD patient group were determined by pairwise comparisons using a non-parametric Wilcoxon rank-sum test. To control for the effect of additional factors, a linear model of the form deltaCT ~ Group + Age + Antidepressant treatment was used. P < 0.05 was considered statistically significant. All the original illustrations used in this manuscript were created with BioRender.com.

## RESULTS

### miR-499-5p expression is increased in human BD patients and the rat social isolation model of early life adversity

Recently, variants in the *MIR499* gene have been associated with BD (20). The main mature miRNA expressed from the MIR499 locus is miR-499-5p, whose function has been extensively studied in the cardiovascular system. To investigate a possible involvement of miR-499-5p in the molecular pathophysiology of BD, we first assessed the levels of miR-499-5p in peripheral blood mononuclear cells (PBMCs) obtained from BD patients (Figure 1A and Supplementary Table 1). Peripheral levels of miR-499-5p were significantly increased in BD patients compared to healthy controls, after correction for age and antidepressant treatment (Figure 1B). Traumatic experiences, especially during childhood, are one of the most important environmental risk factors associated with the development of BD (11, 39). Therefore, for a more comprehensive understanding of the pathogenesis of the disorder, we next asked whether childhood maltreatment influenced the expression of peripheral miR-499-5p. Interestingly, we found that maltreated but otherwise healthy individuals showed a highly significant, almost two-fold increase in miR-499-5p PBMC levels compared to healthy controls without a history of childhood trauma (Figure 1C). Furthermore, the peripheral miR-499-5p levels positively correlated with the severity of maltreatment in general (CTQ Total Score) (Supplementary Figure 1A) and, more specifically, with physical and emotional neglect (Supplementary Figures 1B and C), but not with subscales of abuse (Supplementary Figures 1D, E, and F). Thus, peripheral miR-499-5p expression, possibly as a result of early life adversity, is strongly upregulated in BD patients, further supporting a functional role of miR-499-5p in BD pathogenesis. To determine whether early life adversity also induced the expression of miR-499-5p in the brain, we exposed rats to juvenile social isolation with the aim to model physical and emotional neglect (Figure 1D), which is known to affect behaviors relevant for affective disorders, such as anxiety, drug addiction, and cognitive performance (40–42). As expected, socially isolated rats showed significantly lower hippocampal *c-fos* and *Arc* expression (Figure 1E and F), neural activity-induced genes which are known to be down-regulated upon social isolation in the rat hippocampus (43, 44). Importantly, early life stress induced a highly significant up-regulation (≈3.5 fold) in the expression of miR-499-5p compared to group-housed animals (Figure 1G). The social isolation mediated miRNA upregulation was specific for miR-499-5p, since hippocampal levels of two miRNAs previously implicated in psychiatric disorders (16, 45) were either reduced (miR-146a; Figure 1H) or unaltered (miR-30e; Figure 1I) upon social isolation. Our results show that miR-499-5p regulation by early life stress is conserved between humans and rats, offering an opportunity to study the functional impact of elevated miR-499-5p levels in brain development in the context of an animal model.

**Figure 1:**
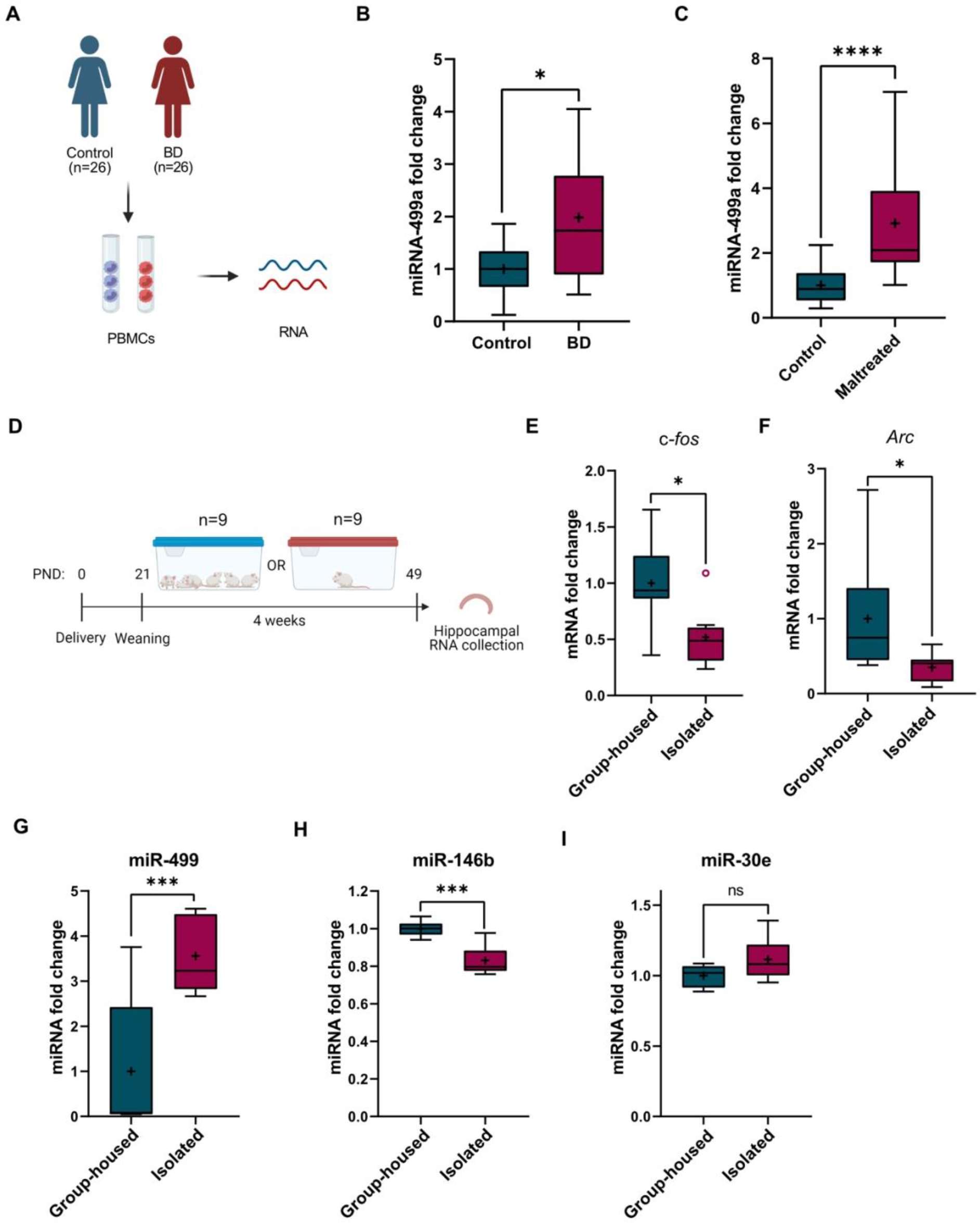
miR-499-5p expression is increased in human BD patients and the rat social isolation model of early life adversity. A) Schematic illustration of the experimental workflow. Total RNA was isolated from PBMCs of psychiatrically healthy subjects (Control, n=25) and BD patients (n=26) for miRNA expression analysis. B) miR-499a qPCR analysis of total RNA isolated from PBMCs of control (n=25) or BD (n=26) subjects (*p=0.0148, Wilcoxon rank-sum test) compared to control subjects after correction for age and antidepressant treatment (linear model of the form delta CT ~ Group + Age + Antidepressant treatment). C) miR-499a qPCR analysis of total RNA isolated from PBMCs of healthy control subjects (Control, n=17) and healthy subjects with a history of childhood maltreatment (Maltreated, n=17). (****p<0.0001, Mann-Whitney U test). D) Schematic representation of the timeline of social isolation rearing. E-I) qPCR analysis for c-fos (E), arc (F), miR-499 (G), miR-146b (H) and miR-30e-5p (I) using total RNA isolated from the hippocampus of male rats that were either group-housed or socially isolated for 4 weeks post-weaning (n=9 rats per group) (*p=0.0207 (c-fos), *p=0.0207 (arc); *** p=0.0008 (miR-499); *** p=0.0006 (miR-146-5p); p=0.1049 (miR-30e-5p); Mann-Whitney U-test). Fold changes represent changes in gene expression relative to the control condition. U6 snRNA was used for normalization. ns = not significant.

### miR-499-5p is expressed in rat hippocampal pyramidal neurons and functions as a negative regulator of dendritogenesis

Since miR-499-5p has not been studied in the mammalian brain, we first measured miR-499-5p levels in different brain regions of the adult rat brain using qPCR. The expression of miR-499-5p was stably detected in all the tested brain regions (Figure 2A), including areas involved in the regulation of mood and cognition, such as the hippocampus and frontal cortex (46, 47). Using single-molecule fluorescence in situ hybridization (smFISH) in dissociated rat primary hippocampal neuron cultures, we detected miR-499-5p positive puncta (red, Fig. 2B) in the soma and, to a lesser extent, dendrites of excitatory pyramidal neurons which co-expressed the marker Camk2a mRNA (green, Fig. 2B). miR-499-5p expression increases during a time-course of hippocampal neuronal development from *DIV* 4 to 20 as assessed by qPCR (Figure 2A), consistent with a function of miR-499-5p in dendrite and/or synapse development. This increase is also observed in glia-depleted cultures (suppl. Fig. 2A), demonstrating the neuronal source of miR-499-5p expression. Given our results obtained from early life adversity models in humans and rats (Fig. 1), we studied a potential regulation of miR-499-5p by stress signaling in rat hippocampal neuron cultures. Glucocorticoids (GCs) are released by the hypothalamic–pituitary–adrenocortical (HPA) axis in response to stressful events and act on neurons via mineralocorticoid (MRs) and glucocorticoid receptors (GRs) to modulate adaptive responses to stress (48, 49). Moreover, high chronic exposure to GCs results in neurotoxicity in models of chronic stress and major depressive disorder (MDD) (50). We observed that a five-day treatment of developing hippocampal neurons with the GR agonist Dexamethasone (DEX) significantly induced the expression of miR-499-5p (Supplementary Figure 2B), which is in line with our previous observations (Figure 1G) and suggests that stress signaling triggers miR-499-5p expression in neurons. Next, we explored potential effects of excessive expression of miR-499-5p, i.e. observed by early life adversity or the activation of stress signaling on neuroplasticity. Therefore, we studied dendritogenesis as a paradigm, since defects in dendritic arborization in the hippocampus are frequently observed in BD patients. To model excessive miR-499-5p activity, we transfected hippocampal neurons with miR-499-5p duplex oligonucleotides (“mimics”), which leads to a strong miR-499-5p overexpression. Sholl analysis revealed that hippocampal neurons transfected with a miR-499-5p mimic displayed a reduced number of intersections compared to control conditions (Figure 2C and D, and Supplementary Figure 2C), indicating reduced dendritic arborization upon miR-499-5p overexpression. Calcium influx through LVGCCs is required for activity-dependent dendritogenesis (10). In addition, both the Ca_v_1.2 pore-forming *Cacna1c* and the regulatory beta-subunit *Cacnb2* are BD risk genes, and impaired cellular calcium homeostasis is a hallmark of BD (51).This led us to hypothesize that Ca_v_1.2 activity could be downstream of miR-499-5p in the control of dendritogenesis. In agreement with this hypothesis, hippocampal pyramidal neurons prepared from constitutive heterozygous *Cacna1c*^+/−^ rats displayed less complex dendritic branches than WT controls (Figure 2E and F), thereby mimicking the effect of miR-499-5p overexpression.

**Figure 2:**
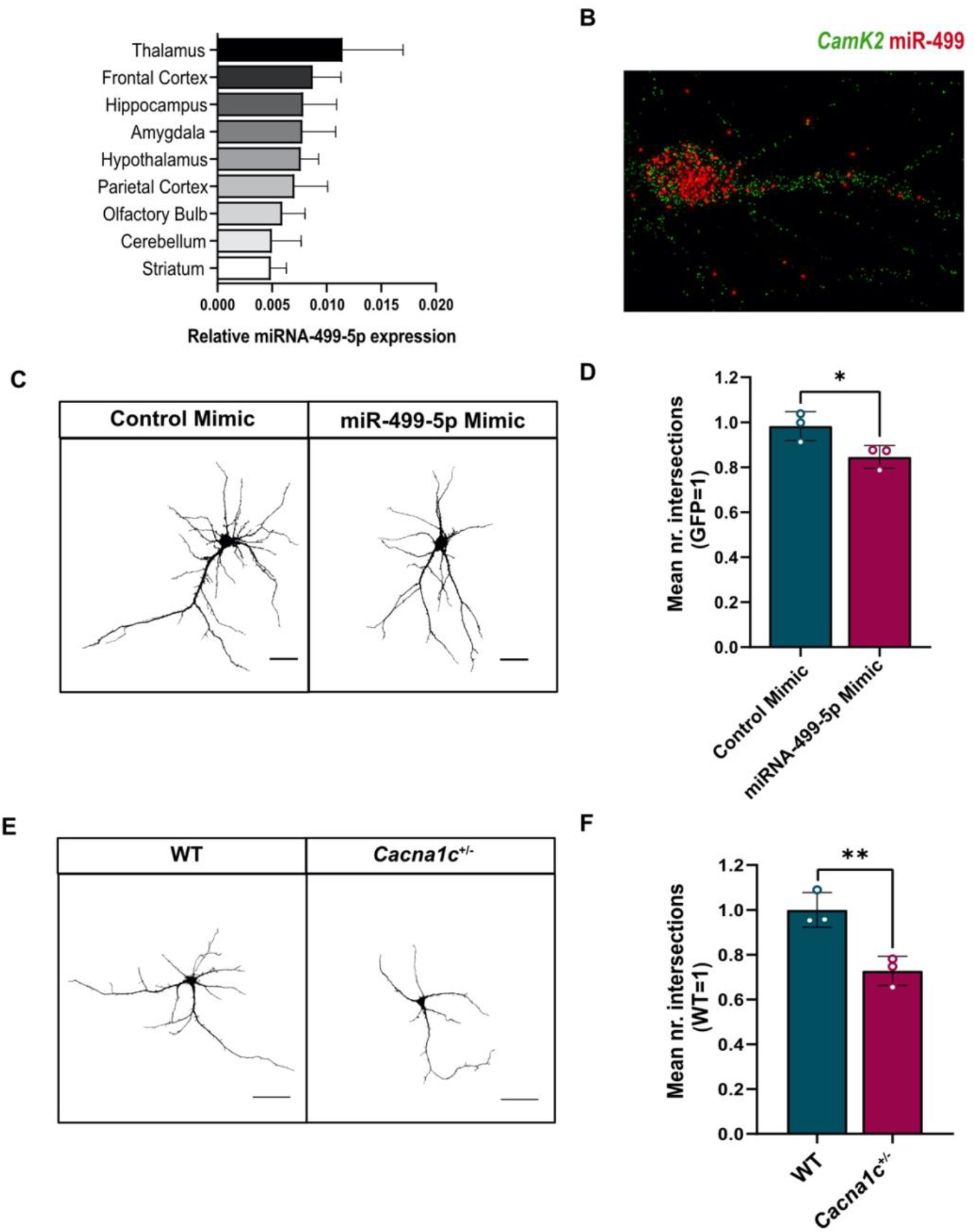
miR-499-5p is expressed in rat hippocampal pyramidal neurons and functions as a negative regulator of dendritogenesis. A) miR-499-5p qPCR analysis using total RNA isolated from different areas of the adult female rat brain (n=3). U6 snRNA was used for normalization. B) Representative picture of single-molecule fluorescence *in situ* hybridization (smFISH) performed in rat hippocampal neurons at DIV7 using probes directed against miR-499-5p (red channel) and CamK2a (green channel) to identify excitatory neurons. C) Representative grey-scale images of primary rat hippocampal neurons (DIV 10) transfected with GFP and the indicated miRNA mimics. Scale bars = 50 μm. D) Quantification of the mean number of intersections by Sholl analysis. GFP-only transfected conditions was set to one in each experiment. (n=3; Unpaired two-sample t-Test, *p=0.0435). E) Representative grey-scale images of GFP-transfected *Cacna1c*^+/+^ (WT) or *Cacna1c*^+/−^ primary rat hippocampal neurons (DIV 10). Scale bars = 50 μm. F) Quantification of the mean number of intersections by Sholl analysis. (n=3; Unpaired two-sample t-Test, **p=0.0096). Fold changes represent changes in dendritic complexity relative to the control condition.

By applying miRNA binding site prediction tools to genes encoding Ca_v_1.2 subunits, we detected a highly conserved miR-499-5p binding site in the 3’UTR of *Cacnb2*, but not *Cacna1c* (Fig. 2G). CACNB2 is a validated target of miR-499-5p in cardiomyocytes (52) and in a very recent GWAS with more than 40 000 BD patients, the CACNB2 locus was significantly associated with BD (53). Therefore, we decided to investigate whether miR-499-5p and CACNB2 functionally interact in rat hippocampal neurons to control dendritogenesis.

### The BD risk gene *Cacbn2* is a downstream target of miR-499-5p in the regulation of hippocampal neuron dendritogenesis

We first studied the effect of miR-499-5p overexpression on Cacnb2 expression. Hippocampal neurons transfected with a miR-499-5p mimic exhibited a significant reduction in the levels of *Cacnb2* mRNA compared to neurons expressing the control mimic as judged by qPCR (Figure 3A). Furthermore, *Cacnb2* mRNA levels were significantly lower in the hippocampus of socially isolated rats compared to group-housed rats (Figure 3B). The direction of change in socially isolated rats is opposite to the one observed for miR-499-5p (Fig. 1G), consistent with a negative regulatory role for miR-499-5p in Cacnb2 expression. To test a possible physical interaction of miR-499-5p with the *Cacnb2* 3’UTR, we performed reporter gene assays in primary hippocampal neurons using luciferase genes fused to either the wild-type (WT) Cacnb2 3’UTR or Cacnb2 3’UTR containing mutations in the predicted miR-499-5p seed match (MUT) (Figure 3C). Overexpression of miR-499-5p resulted in a significant repression of luciferase reporter gene expression for the *Cacnb2* WT plasmid (Figure 3D), consistent with a post-transcriptional, 3’UTR-dependent repressive effect of miR-499-5p on Cacnb2. This effect was not observed when the *Cacnb2* MUT plasmid was transfected (Figure 3D), demonstrating that miR-499-5p mediated its repressive effect via direct interaction with the seed match site in the *Cacnb2* 3’UTR. On the other hand, the inhibition of endogenous miR-499-5p in neurons by transfection of an LNA-modified antisense oligonucleotide (miR-499-5p pLNA) led to a significant increase in the expression of the *Cacnb2* WT, but not Cacnb2 MUT, luciferase reporter gene (Figure 3E). Together, these experiments establish Cacnb2 as a *bona-fide* miR-499-5p target gene in rat hippocampal neurons. We next investigated whether miR-499-5p mediated regulation of Cacnb2 is functionally involved in dendritogenesis. Therefore, we transfected hippocampal neurons with a miR-499-5p mimic and examined whether restoring *CACNB2* expression by co-transfection of a CACNB2 expression plasmid (pMT2-CACNB2; Supplementary Figure 5) was able to rescue impaired dendritogenesis observed upon miR-499-5p overexpression. Notably, whereas overexpression of miR-499-5p reduced dendritic complexity as expected, dendritic complexity of neurons co-transfected with miR-499-5p and pMT2-CACNB2 was not significantly different compared to control conditions (Figure 3F and G). This result provides strong evidence that *Cacnb2* is an important downstream component of the miR-499-5p-mediated regulation of dendritic development.

**Figure 3:**
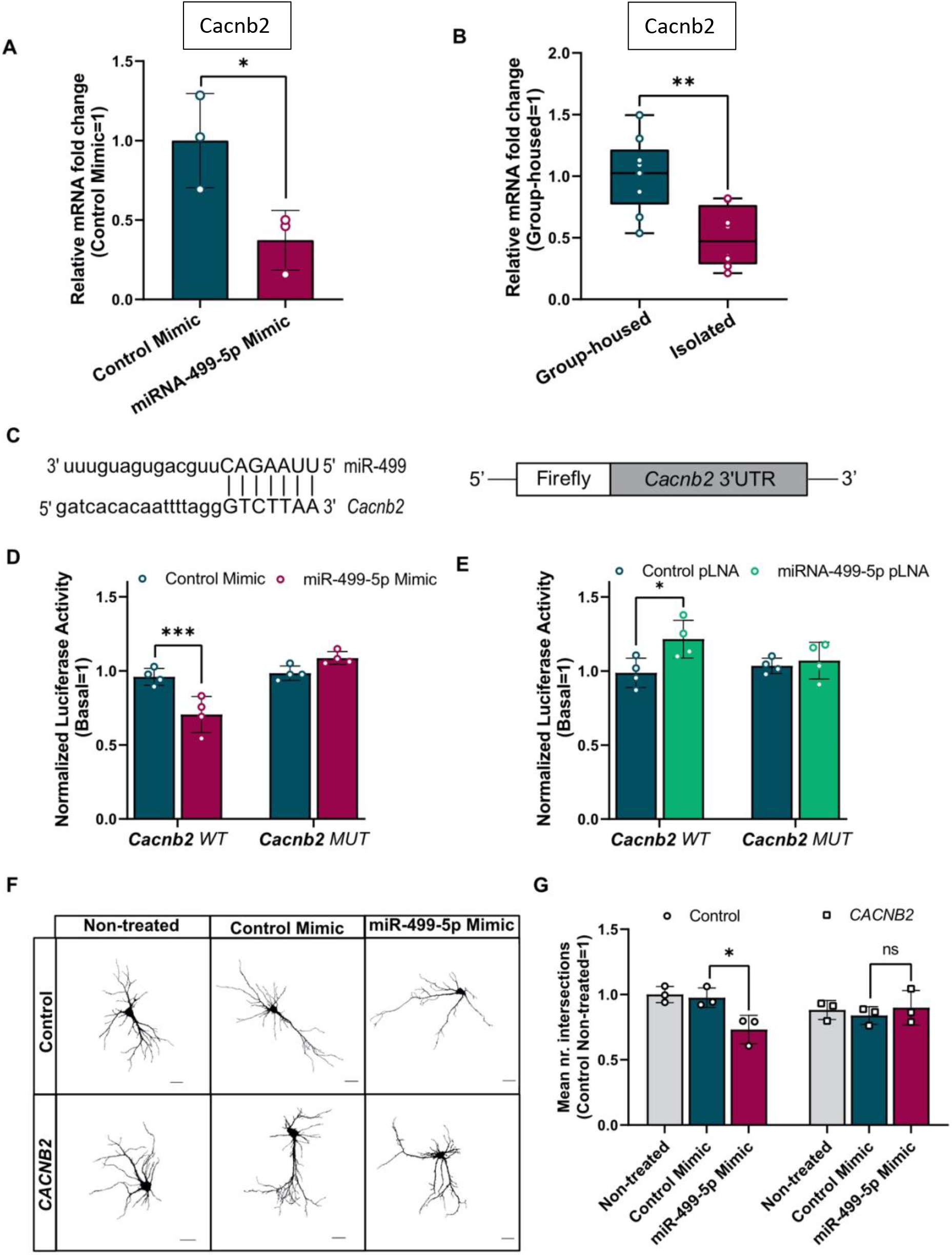
The BD risk gene *Cacbn2* is a downstream target of miR-499-5p in the regulation of hippocampal neuron dendritogenesis. A) qPCR analysis of *Cacnb2* mRNA levels using total RNA isolated from rat hippocampal neurons transfected with either a control or miR-499-5p mimic (n=3; Unpaired two-sample t-test, *p=0.0363). B) qPCR analysis of *Cacnb2* mRNA levels using total RNA from the hippocampus of either socially isolated or group-housed rats (n=9 rats per group; Unpaired two-sample t-test, **p=0.0020). C) Nucleotide base pairing of miR-499-5p with *Cacnb2* 3’UTR (left) and schematic representation of the *Cacnb2* luciferase reporter (right). D) Relative luciferase activity of rat hippocampal neurons transfected with the indicated miRNA mimics and expressing either a *Cacnb2* WT or MUT reporter (n=4; Two-way ANOVA: main effect of the miRNA mimic p=0.0087, of the *Cacnb2* luciferase reporter p<0.0001 and of the miRNA mimic by Cacnb2 luciferase reporter interaction p<0.0001, Post hoc Tukey test: ***p=0.0002). E) Relative luciferase activity of rat hippocampal neurons transfected with the indicated pLNAs and expressing either a *Cacnb2* WT or MUT reporter (n=4; Two-way ANOVA: main effect of the miRNA pLNA p=0.0061, no main effect of the *Cacnb2* luciferase reporter p=0.3590 or the miRNA pLNA by *Cacnb2* luciferase reporter p=0.0910, Post hoc Tukey test: *p=0.0152). Fold changes represent changes in relative luciferase activity of transfected neurons relative to the non-transfected neurons. F) Representative images of *DIV* 10 hippocampal neurons co-transfected with control or miR-499-5p mimics and a pMT2-*CACNB2* expression plasmid (*CACNB2*). Scale bars = 50 μm. G) Quantification of the mean number of intersections by Sholl analysis. (n=3; Two-way ANOVA: no main effect of the *CACNB2* transfection p=0.5027 or the miRNA mimics p=0.0805, main effect of the *CACBN2* transfection by miRNA mimics interaction p=0.0213, Post hoc Tukey test: *p=0.0303). Fold changes represent the changes in dendritic complexity of transfected neurons compared to non-treated control neurons. ns = not significant.

### Elevated levels of miR-499-5p in hippocampal neurons impair cell surface expression and activity of Ca_v_1.2 channels

The auxiliary β_2_ subunit of LVGCCs is an essential regulator of the channel cell surface expression (54). To assess whether the regulation of *Cacnb2 by miR-499-5p* had an effect on the surface expression of Ca_v_1.2 channels, we performed anti-HA immunostaining on non-permeabilized hippocampal neurons (“live staining”) which had been transfected with an HA-tagged α_1_ pore-forming subunit (Cacna1c-HA) (55) together with miR-499-5p mimic. Since the Cacna1c-HA-epitope is only recognized once exposed on the cell surface, this procedure allows to measure Ca_v_1.2 cell surface expression. In accordance with previous work showing a post-synaptic localization of Cav1.2 at dendritic shafts and cell bodies (56), we detected Cav1.2-HA puncta in both compartments (Figure 4A). However, surface expression of Ca_v_1.2-HA was reduced upon overexpression of miR-499-5p as quantified by a significant decrease in the cell area covered by cell surface clusters of Cav1.2-HA channels (Figure 4B), integrated density, and Ca_v_1.2-HA area (Supplementary Figures 6A and B). We did not observe any changes in cell area covered by total Ca_v_1.2-HA (Figure 4C), integrated density, or total Cav1.2 area (Supplementary Figures 4C and D) when we performed the HA staining under permeabilized conditions, suggesting that miR-499-5p overexpression did not affect overall levels of recombinant Ha-tagged *Cacna1c*. Since Ca_v_1.2 is the predominant LVGCC isoform in hippocampal neurons (54), we hypothesized that the observed reduction in cell surface expression of Ca_v_1.2 channels translated into a corresponding loss of LVGCC currents (I_Ca,L_). To test this hypothesis, whole-cell patch-clamp recordings were performed on hippocampal neurons overexpressing miR-499-5p. Calcium currents were evoked in conditions that allowed isolation of I_Ca,L_ as indicated by a ≈50% reduction in current density by nifedipine treatment (Supplementary Figure 4E). Consistent with our results from immunostaining (Fig. 4A–C), hippocampal neurons overexpressing miR-499-5p showed significantly smaller LVGCC current density than control neurons (Figure 4D and E). Neither channel activation (Supplementary Figure 4F) nor inactivation (Supplementary Figure 4G) were significantly altered by miR-499-5p overexpression, suggesting that miR-499-5p affected LVGCC activity primarily by regulating Ca_v_1.2 membrane incorporation. Furthermore, transfection of the miR-499-5p mimic led to a decrease in membrane capacitance, which however did not reach statistical significance (Supplementary Figure 4H). This finding correlates well with reduced dendrite arborization observed upon miR-499-5p overexpression, which is supposed to lead to an overall decrease in neuronal membrane area (Figure 2C and D). We conclude that miR-499-5p negatively regulates the function of Ca_v_1.2 calcium channels most likely by reducing their surface expression due to an inhibition of Cacnb2 expression.

**Figure 4:**
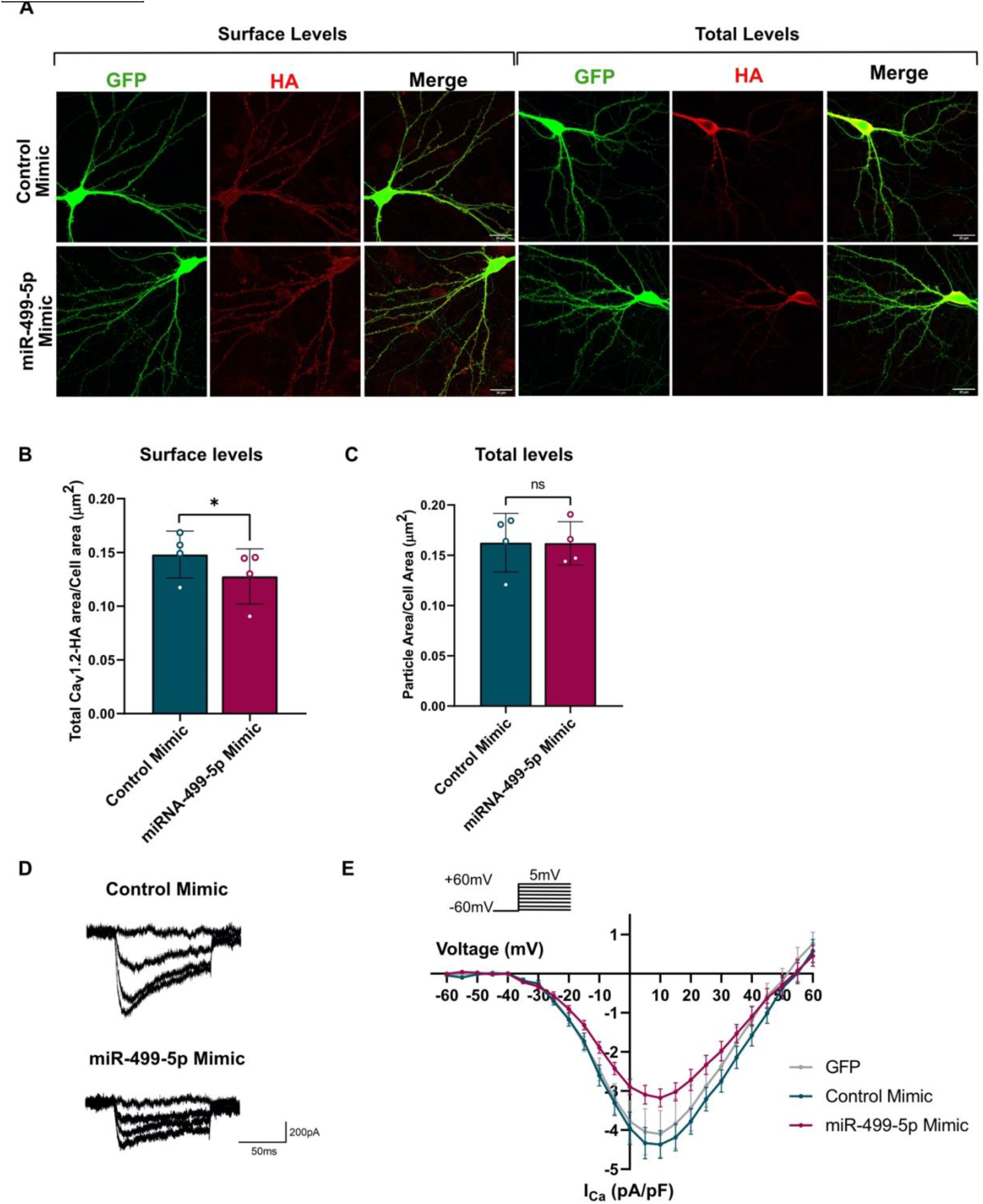
Elevated levels of miR-499-5p in hippocampal neurons impair cell surface expression and activity of Ca_v_1.2 channels. A) Representative images of *DIV* 19 rat hippocampal neurons co-transfected with GFP (green channel) and Cav1.2-HA (red channel), together with either control or miR-499-5p mimics. After 12-13 days of expression, labeling with Anti-HA antibodies was performed in live conditions to identify cell surface Cav1.2 channels (left) or under permeabilized conditions to identify total levels of Cav1.2 channels (right). Scale bars = 20 μm. B) Quantification of the area occupied by surface Cav1.2 fluorescence normalized to the total cell area (GFP fluorescence) from non-permeabilized neurons transfected as in A) (n=4; Paired two-sample t-Test *p=0.0318). C) Quantification of the total levels of Cav1.2 channels of permeabilized neurons transfected as in A). D) Representative calcium current traces from neurons transfected with the indicated miRNA mimics. E) I/V curves (current density against voltage) of calcium peak currents from hippocampal neurons co-transfected with GFP (n=9 cells) and the Control (n=10 cells) or miR-499-5p mimic (n=10 cells) (p<0.05 for miR-499-5p mimic vs. Control mimic between −10 and +30 mV, Unpaired two-sample t-Test).

### miR-499-5p represses CACNB2 expression in the rat hippocampus *in vivo* and impairs short-term recognition memory in *Cacna1c*^+/−^ rats

To further explore the impact of miR-499-5p dysregulation and Ca_v_1.2 calcium channel dysfunction *in vivo*, we decided to overexpress miR-499-5p in the rat hippocampus. Therefore, a recombinant adeno-associated virus (rAAV) expressing a miR-499-5p precursor RNA together with eGFP under the control of the human synapsin promoter (AAV-miR-499) was bilaterally injected into the rat dorsal and ventral hippocampus of adult rats (Figure 5A; suppl. Fig. 5). rAAV-mediated miR-499-5p delivery led to significant overexpression of miR-499-5p (Fig. 5B; suppl. Fig. 6A, B). CACNB2 protein levels, as assessed by Western blot, were significantly reduced in the hippocampus of rats overexpressing miR-499-5p and negatively correlated with miR-499-5p levels (Fig. 5C, D; suppl. Fig. 7). These findings confirm the negative regulation of CACNB2 by miR-499-5p in the rat brain *in vivo*.

**Figure 5:**
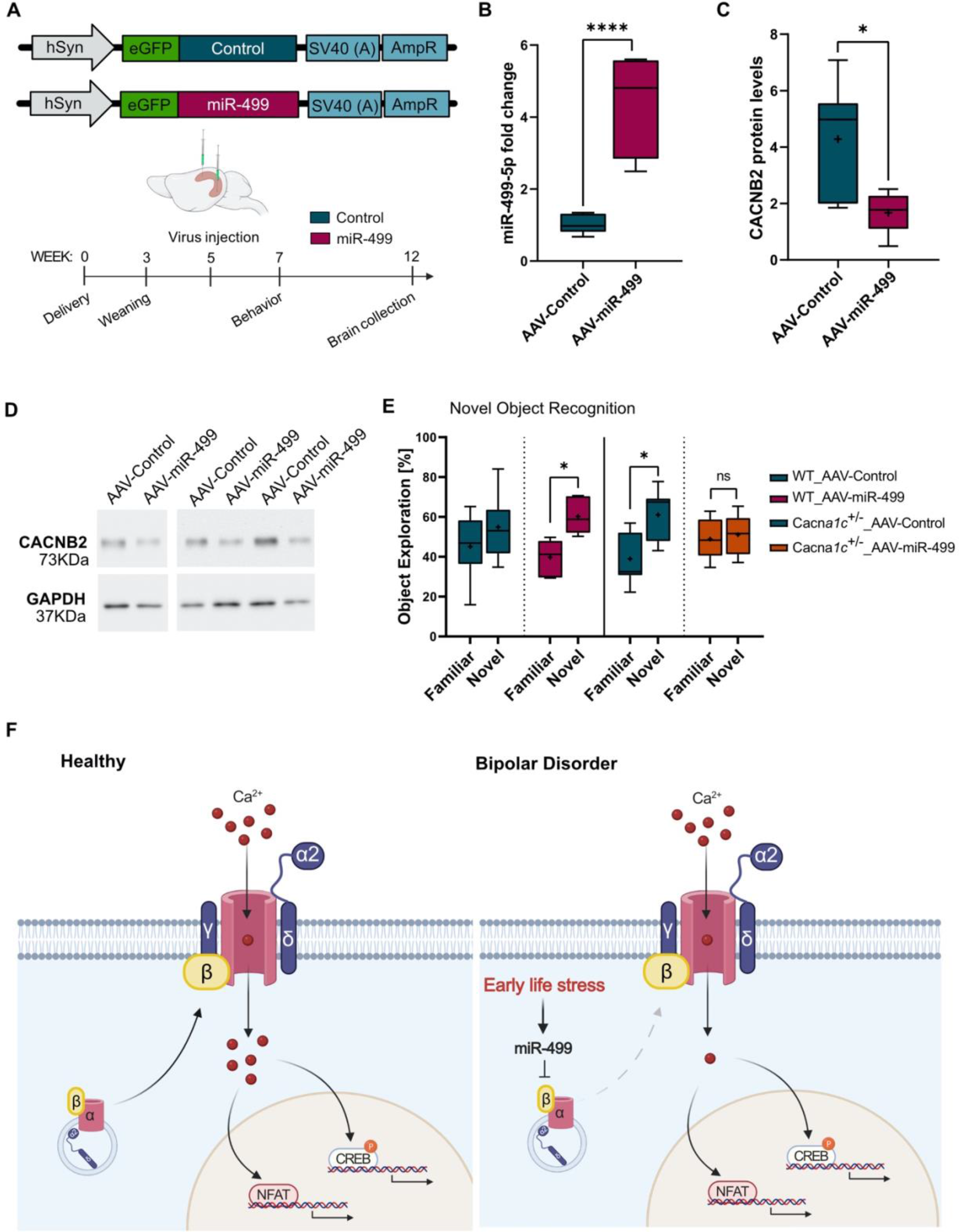
miR-499-5p represses CACNB2 expression in the rat hippocampus *in vivo* and impairs short-term recognition memory in *Cacna1c*^+/−^ rats. A) Schematic illustration of the hSyn-eGFP-scramble control (AAV-Control) and hSyn-eGFP-chimeric miR-499-5p hairpin (AAV-miR-499), and timeline of virus infection experiments. B) miR-499-5p qPCR analysis from total RNA isolated from the right hippocampus of rats injected with either AAV-control or AAV-miR-499 (****p<0.0001, Unpaired two-sample t-Test; n=7 rats in the AAV-Control group, n=6 rats in the AAV-miR-499 group). C) Quantification of CACNB2 Western blots using hippocampal protein lysate from rats injected with either AAV-control or AAV-miR-499 (*p=0.0103, Unpaired two-sample t-Test; n=7 rats in the AAV-Control group, n=6 rats in the AAV-miR-499 group). D) Representative Western blot images of *CACNB2* (upper panel) and *GAPDH* (lower panel) protein expression levels in the hippocampus of WT rats injected with the AAV-Control or AAV-miR-499 hairpin viruses. GAPDH was used as a loading control. E) Novel object recognition task. Percentage of time WT or *Cacna1c+/-* rats injected with the indicated AAV explored either the familiar or novel object. Data are presented as means ±SEM (*p<0.05, paired two-sample t-Test; WT_Control: n=12; WT_miR-499: n=6; *Cacna1c*^+/−^_Control: n=9; *Cacna1c*^+/−^_miR-499: n=8). F) Proposed model for the mechanism of miR-499-5p dysregulation in BD. Early life stress induces the expression of miR-499-5p in human PBMCs and rat hippocampus, which impairs dendritic development and Ca_v_1.2 calcium channel cell surface expression and activity by inhibiting the expression of an auxiliary subunit of Ca_v_1.2 calcium channels and risk gene for BD, CACNB2.

We went on to investigate a potential role of elevated miR-499-5p for behavioural phenotypes associated with BD. For these experiments, we initially focused on the assessment of cognitive function, since LVGCC activity was repeatedly shown to affect memory-related functions (7). In the novel object recognition test, which is affected by hippocampal lesions (57), rats typically spent more time exploring the novel object than the familiar one as measured by the increase in the time spent sniffing (percentage of total exploration) the novel object compared to the familiar object (Figure 5E). Overexpression of miR-499-5p in the hippocampus of WT animals did not alter the animal’s preference for the novel object as the animals spent significantly more time exploring the novel object. However, in *Cacna1c*^+/−^ rats overexpressing miR-499-5p the preference for the novel object was almost completely lost (Figure 5E). Thus, miR-499-5p overexpression selectively impairs short-term recognition memory in the context of reduced Ca_v_1.2 expression.

## DISCUSSION

The pathophysiology of BD is highly complex, involving the interaction between several genetic modifications and environmental insults with functional and structural effects that are poorly understood and difficult to disentangle. More than 50% of patients report having experienced childhood maltreatment (58). A recent meta-analysis of 30 independent studies found that BD patients with a history of childhood maltreatment present more extreme manic, depressive and psychotic symptoms, higher risk of comorbidity with other psychiatric illnesses, early disease onset, risk of rapid cycling, and higher risk of suicide attempt (11). Importantly, maltreated patients respond differently to treatment compared to patients without childhood trauma (59). In this study, we showed that miR-499-5p is increased in PBMCs of BD patients and healthy individuals with a history of childhood maltreatment, in particular physical and emotional neglect. By systematically varying early life experiences in rats, we further showed that juvenile social isolation, mimicking aspects of physical and emotional neglect, leads to an upregulation of miR-499-5p in the brain. Thus, our results point towards an important role of miR-499-5p upregulation as an early marker of childhood trauma and unfavorable course of illness.

The observation that early life stress induced the expression of miR-499-5p in the blood of maltreated subjects and the hippocampus of socially isolated rats raises the question of the source of miR-499-5p overexpression. MiR-499-5p is an intronic miRNA encoded by the Myosin heavy chain 7B (Myh7b) gene with particularly high expression in the cardiovascular system (21). Thus, chronic stress might lead to aberrant miRNA-499-5p expression in the heart and secretion of miR-499-5p into the bloodstream inside exosomes that then reach the brain. In agreement with this hypothesis, dysregulated expression of miR-499-5p in prefrontal cortex (PFC) exosomes was previously reported in schizophrenia (SZ) and BD patients (60). Exosomal transfer of miR-499-5p between the cardiovascular system and the brain would be especially relevant in light of the high co-morbidity between heart disease and BD (39). Indeed, higher levels of circulating miR-499-5p were reported in patients of acute myocardial infarction (61), cardiomyopathy (62), and atrial fibrillation (52). Mechanistically, activation of the MYH7B/MIR499 gene upon early life stress could involve epigenetic modifications, since Tsumagari et al. showed that exons 10, 17, and 18 of the MYH7B/MIR499 gene were hypomethylated in heart tissue compared to muscle progenitor cells, which correlated with higher miR-499-5p levels (63).

Our result that the negative effect of miR-499-5p overexpression on dendritogenesis was rescued by co-expression of *CACNB2* strongly implicates LVGCC activity as an important downstream component of miR-499-dependent regulation of neuroplasticity. Previous studies already demonstrated an important role of LVGCC signaling in dendritogenesis in the context of neuropsychiatric disease. For example, inhibition of LVGCCs with Nifedipine significantly reduced dendritic growth (64), and activation of the Ca_v_1.2 downstream effectors CaMK, MAPK, and CREB are critical for dendritic growth (10). Interestingly, iPSC-derived neurons from Timothy syndrome (TS) patients, which carry a gain-of-function mutation in the CACNA1C gene, are prone to activity-dependent dendrite retraction (65). Intriguingly, this phenotype does not depend on calcium influx through Ca_v_1.2, but rather on the recruitment of the small Rho-family GTPAse Gem to Ca_v_1.2 (66). By analogy, the reduction in *CACNB2* may impair dendritic growth by destabilizing the interaction between Gem and Ca_v_1.2 calcium channels. In addition, the β_2_ subunit is necessary for the trafficking of fully matured channel complexes from their site of folding to the cell membrane. Consistent with these observations, overexpression of miR-499-5p in hippocampal neurons led to reduced Ca_v_1.2 cell surface expression and current density. In HEK293 cells, Cacnb2 increased the surface levels of Cav1.2 channels by preventing channel ubiquitination by the E3 ubiquitin ligase Ret Finger Protein 2 (RFP2) and entry into the ER-associated protein degradation system (67). Therefore, in addition to effects on forward trafficking, miR-499-5p overexpression might lead to increased endocytosis followed by proteasome-mediated degradation. Further studies are warranted to obtain more conclusive results regarding the mechanism of action of miR-499-5p.

Although an imbalance in cellular calcium homeostasis is widely accepted in the BD field, it is still controversial in which direction calcium concentrations are actually changed. Intracellular calcium levels are overall increased in the plasma and lymphocytes of BD patients but decreased in olfactory neurons. In human iPSC-derived neurons from individuals carrying the *CACNA1C* risk allele rs1006737, higher mRNA levels of *CACNA1C* and higher L-type calcium currents compared to neurons carrying the non-risk variant are observed (68). However, the *CACNA1C* risk allele rs1006737 has been shown to result in both gain- and loss-of-function of *CACNA1C* gene expression (69, 70), depending on the brain region examined. Deletion of *Cacna1c* in mice induces the expression of anxiety-like behaviors but has an anti-depressant effect (71), while in rats it was shown to impair social behavior (27). Importantly, the effect of *Cacna1c* haploinsufficiency seems to depend on the developmental stage. Embryonic deletion of *Cacna1c* in forebrain glutamatergic neurons resulted in increased chronic stress susceptibility while during adulthood produced stress resilience (72). Taken together, these observations suggest that calcium signaling alterations in both directions are possibly detrimental and can lead to defects in neuroplasticity, e.g. dendritic impairments, depending on the brain region, developmental stage, and the history of chronic stress.

A few previous studies already indicated that alterations in miR-499-5p expression could contribute to the development of psychiatric disorders. For example, miR-499 is downregulated In exosomes derived from the PFC of SZ and BD patients (60). On the other hand, *post mortem* PFC of depressed subjects shows higher levels of miR-499-5p (73), which would be more in line with our observation of miR-499-5p upregulation in PBMCs of subjects suffering from early life adversity or BD. These inconsistencies could reflect again the fact that miR-499 levels have to be within a physiological window, and that deviations in both directions might be detrimental. According to this hypothesis, a physiological induction of miR-499-5p by stress might in fact be necessary to maintain calcium homeostasis and prevent the development of BD. Interestingly, this view is supported by our unpublished observation that a BD-associated mutation in the miR-499 gene prevents the production of mature miR-499 by impairing Drosha-mediated pri-miR-499 cleavage (Verhaerts, Martins, et al., unpublished). Since miRNAs usually have multiple target mRNAs, other miR-499-5p targets should also be considered. One interesting candidate is the validated miR-499-5p target PDCD4, an endogenous inhibitor of BDNF translation. *Pdcd4* expression is induced by chronic stress, and PDCD4 overexpression in the mouse hippocampus triggers depression-like behaviors (74). Genome-wide expression profiling should help to identify additional miR-499-5p targets which can be followed-up in functional experiments.

Cognitive impairments, especially related to working memory and executive functioning, are among the core symptoms of BD in addition to mood disturbances (1). Our finding that overexpression of miR-499-5p in the hippocampus of *Cacna1c*^+/−^ rats impairs short-term recognition memory argues in favor of an important contribution of miR-499 dysregulation to cognitive symptoms associated with BD. In the novel object recognition test, neither the genotype nor miR-499-5p overexpression alone affected the preference for the novel object compared to the familiar one. This observation agrees with a previous study which reported a normal preference for the novel object in *Cacna1c*^+/−^ and WT rats of both sexes (38). Apparently, miR-499-5p mediated reduction in calcium signaling is not sufficient to elicit LTP and associated memory impairments. In contrast, in *Cacna1c*^+/−^ rats, overexpression of miR-499-5p reduces the surface availability and activity of Ca_v_1.2 channels to an extent where memory formation is no longer supported. *Cacna1c*^+/−^ mice also showed deficits in spatial memory (7) and extinction of contextual fear (75). Further behavioral tests are required to obtain a more comprehensive picture of cognitive abilities, as well as phenotypes related to mania and depression, of rats overexpressing miR-499-5p.

In conclusion, we describe a novel mechanism for calcium signaling dysfunction in BD involving miRNA-dependent regulation of the BD risk gene *Cacnb2*. Early life adversity-induced miR-499-5p represses *CACNB2* translation and reduces calcium influx required for dendritic development and cognitive function. Furthermore, circulating miR-499-5p could be used as a biomarker for the stratification of BD patients based on their history of childhood trauma or to identify healthy individuals that are at elevated risk to develop a psychiatric condition and would therefore benefit from preventative therapies. Finally, miR-499-5p inhibition, e.g. via stable brain delivery of a chemically modified, LNA-based miR-499-5p antisense oligonucleotide, could represent a new avenue for the treatment of cognitive impairments associated with neuropsychiatric disorders.

## ACKNOWLEDGMENT

We acknowledge the excellent technical assistance provided by Tatjana Wüst and Christina Furler. David Colameo wrote a Python script for Image Analysis. We are further grateful for the kind gift of the HA-Cacna1c expression plasmid by Amy Lee (Iowa). The authors wish to thank Christine Hohmeyer, Marcella Rietschel, and Stephanie H. Witt for their help with genotyping. This work was supported by grants from the Swiss National Foundation (SNF 310030E_179651, 32NE30_189486) to G.S. This work is further part of the Forschergruppe “Neurobiology of affective disorders: Translational perspectives on brain structure and function” (FOR 2107) and was supported by grants from the Deutsche Forschungsgemeinschaft to R.S. (DFG 559/14-1 and DFG 559/14-2), to M.W. (DFG WO 1732/4-1 and DFG WO 1732/4-2) and to T.K. (DFG KI 588/14-1 and KI 588/14-2).

## AUTHOR CONTRIBUTIONS

HCM performed RNA isolation from PBMCs and rat tissue, qPCR, dendritogenesis assays, Western blots, luciferase reporter gene assays, immunostaining, plasmid cloning and wrote the manuscript. AÖS performed stereotactic injections into the rat hippocampus and behavioural experiments with the help of NMR. CG performed histology and qPCR. MP helped with RNA isolation from PBMCs and qPCR. SB performed smFISH experiments. FG performed statistical analysis of human miRNA expression data. JW performed electrophysiological recordings. MDB and TMK performed the juvenile social isolation experiments in *Cacna1c* haploinsifficient rats. MW conceptualized and supervised the study on stereotactic injections into the rat hippocampus and behavioural experiments and the juvenile social isolation experiments in *Cacna1c* haploinsifficient rats. GS conceptualized and supervised the project, wrote the manuscript and coordinated collaborative experiments. MW helped revising the manuscript. GS, RKWS, and MW acquired funding and provided resources. TK conceptualized the overall project, obtained funding, supervised human data collection, and commented the manuscript.

---

**Supplementary Table 1:** Subjects used for miR-499-5p expression analysis on psychiatrically healthy controls (Control) or Bipolar disorder patients (BD). One-way-ANOVA was performed to evaluate significant differences between groups. s.d.: standard deviation, CTQ: Childhood Maltreatment Questionnaire, HAMD: Hamilton Depression Rating Scale, YMRS: Young Mania Rating Scale, AD: antidepressant use.

|  | **Control** | **BD** | **P value** |
| --- | --- | --- | --- |
| *n* | 26 | 26 | N/A |
| Sex | Female | Female | N/A |
| Age ± s.d. | 29.4 ± 5.6 | 29.9 ± 5.5 | 0.94 |
| CTQ ± s.d. (%) | 14 ± 11.4 (53.8%) | 8 ± 20.2 (53.8%) | 0.12 |
| Family history of BD (%) | 8 (30.8%) | 11 (30.8%) | N/A |
| HAMD ± s.d. | 2.7 ± 3.9 | 7.5 ± 6.2 | <0.0001 |
| YMRS ± s.d. | 0.5 ± 1.1 | 2.4 ± 2.9 | <0.01 |
| AD (%) | 0 (0%) | 7 (26.9%) | N/A |

**Supplementary Table 2:**
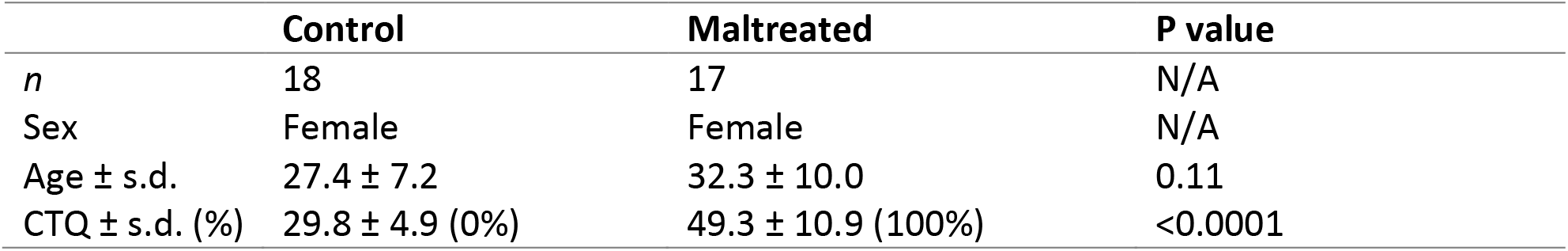
Subjects used for miR-499-5p expression analysis in healthy controls (Control) and subjects with a history of childhood maltreatment (Maltreated). Unpaired t-Test was performed to evaluate significant differences between groups. s.d.: standard deviation, CTQ: Childhood Maltreatment Questionnaire, AD: antidepressant use.

|  | **Control** | **Maltreated** | **P value** |
| --- | --- | --- | --- |
| *n* | 18 | 17 | N/A |
| Sex | Female | Female | N/A |
| Age ± s.d. | 27.4 ± 7.2 | 32.3 ± 10.0 | 0.11 |
| CTQ ± s.d. (%) | 29.8 ± 4.9 (0%) | 49.3 ± 10.9 (100%) | <0.0001 |

**Supplementary Table 3:**
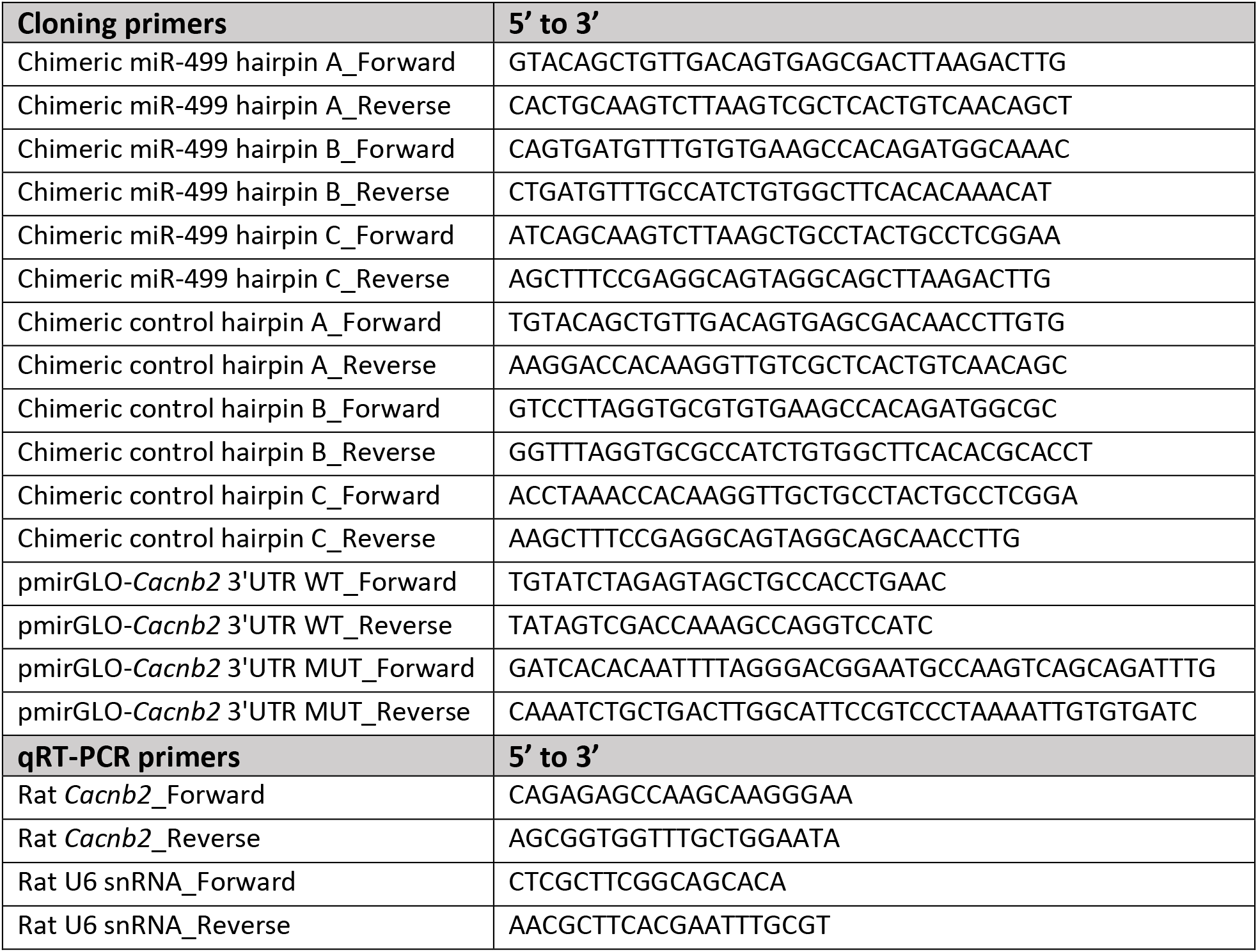
Primer sequences used for cloning and RT-PCR.

**Supplementary Figure 1:**
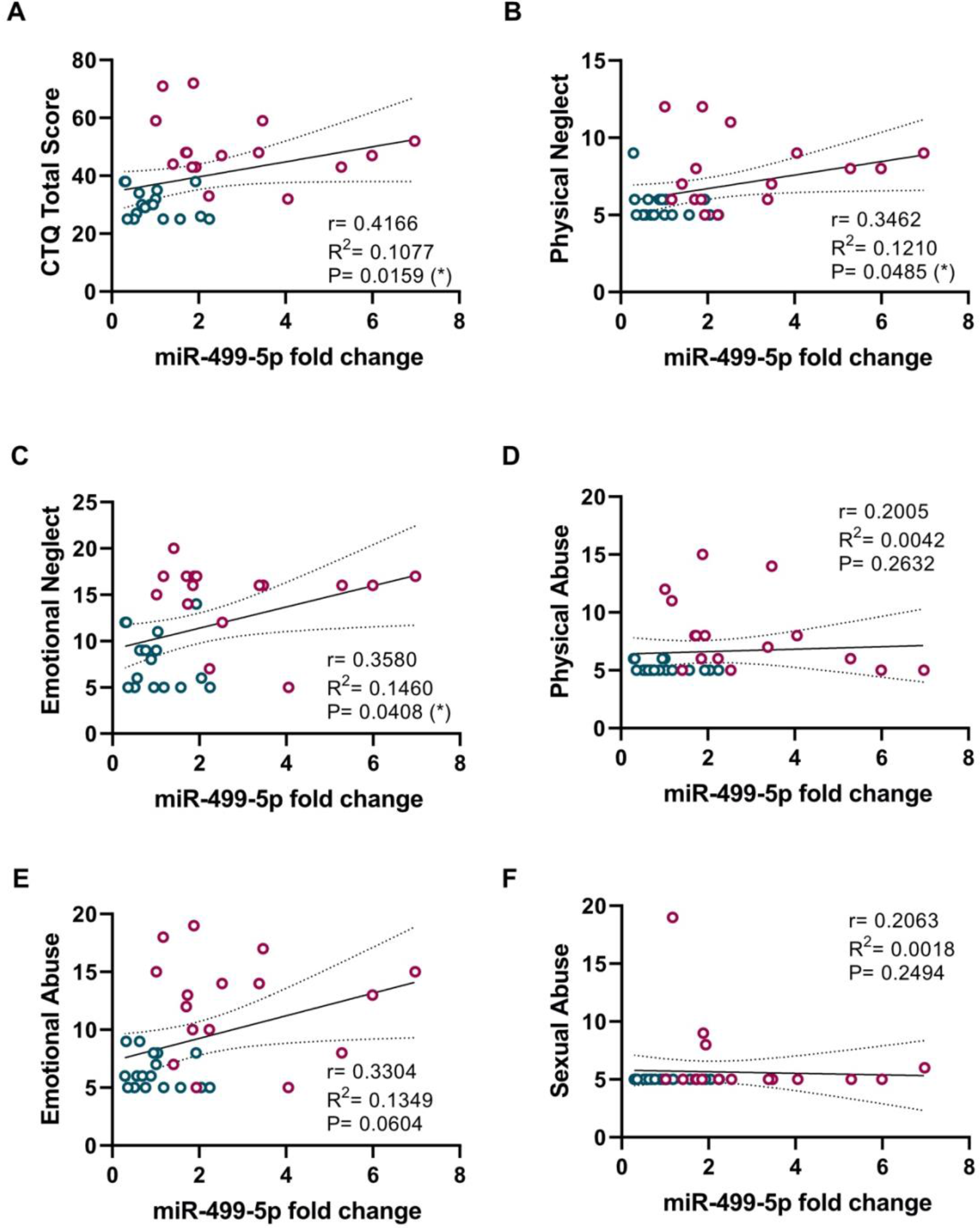
Positive correlation between the peripheral levels of miR-499-5p and A) the sum of all subscale CTQ scores (CTQ Total Score), B) the scores for physical neglect, and C) the scores for emotional neglect. No correlation was found between the peripheral levels of miR-499-5p and the scores for D) physical abuse, E) emotional abuse and F) sexual abuse. Fold changes represent changes in maltreated subjects relative to healthy controls.

**Supplementary Figure 2:**
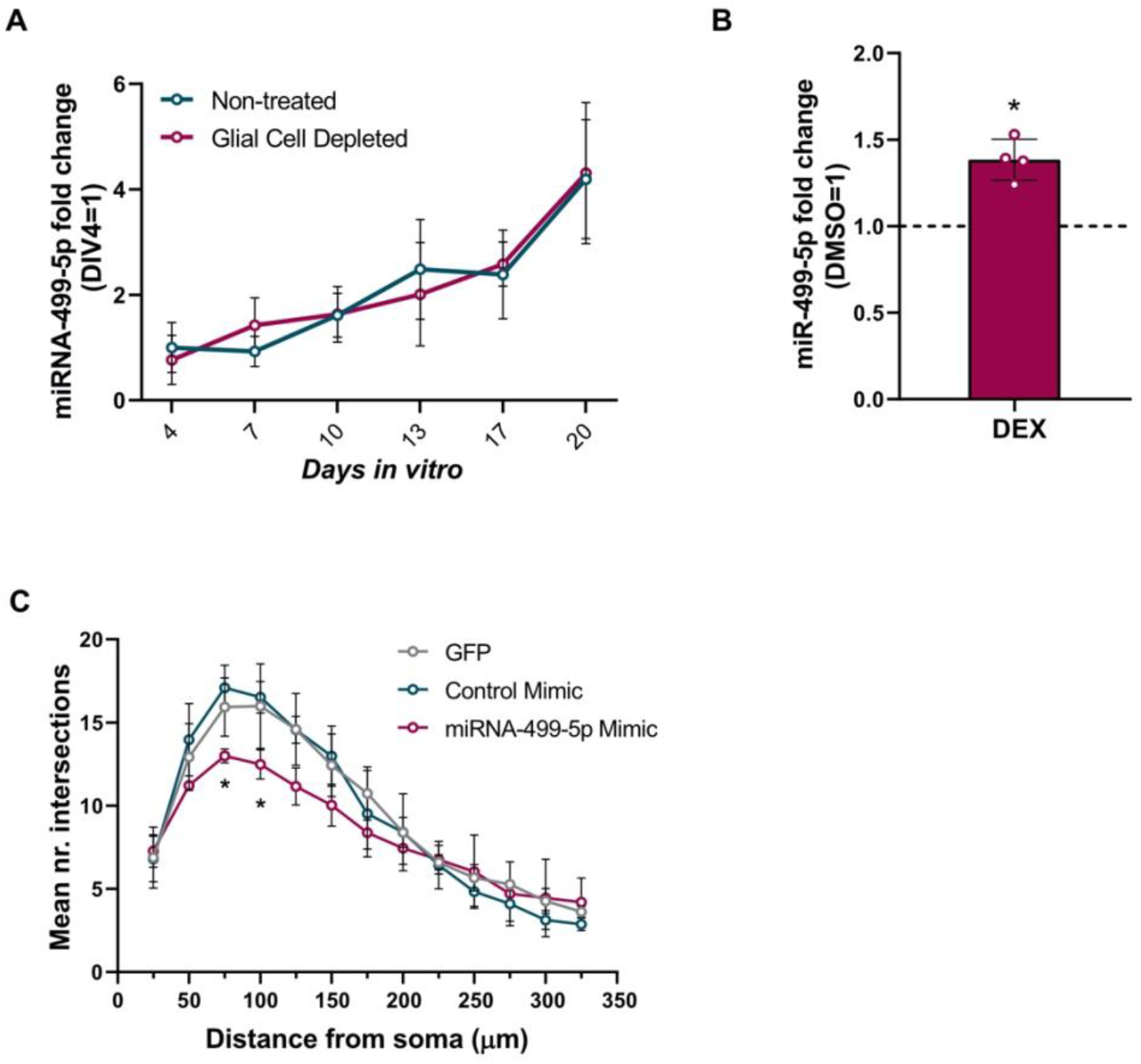
A) Relative expression of miR-499-5p in primary hippocampal neurons at different *DIVs* (n=3). Total RNA was obtained from non-treated developing hippocampal neurons or treated from DIV 3 with FUDR to stop the proliferation of non-neuronal glial cells at the six indicated time points. Fold changes represent changes in miR-499-5p expression relative to DIV 4. B) Relative expression of miR-499-5p is significantly induced in hippocampal neurons treated with DEX compared to DMSO-treated neurons (n=4; One-sample t-Test, **p=0.0075). Fold change represents changes in miR-499-5p expression of DEX-treated neurons relative to DMSO-treated neurons. C) Mean of the Sholl profile averages from biological replicates of Fig. 2C and D.

**Supplementary Figure 3:**
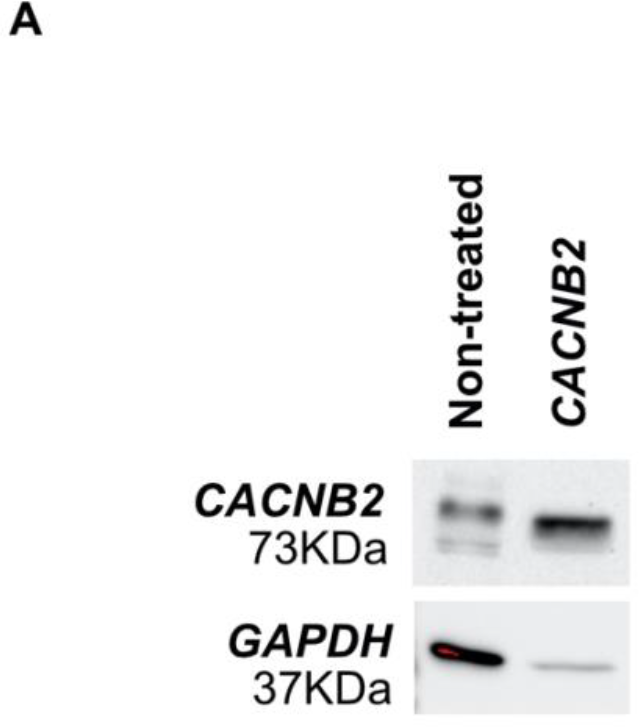
Representative western blot image of *CACNB2* and *GAPDH* protein expression levels. Transfection of cortical neurons with the *CACNB2* expressing construct overexpressed *CACNB2* protein compared to non-transfected cells.

**Supplementary Figure 4:**
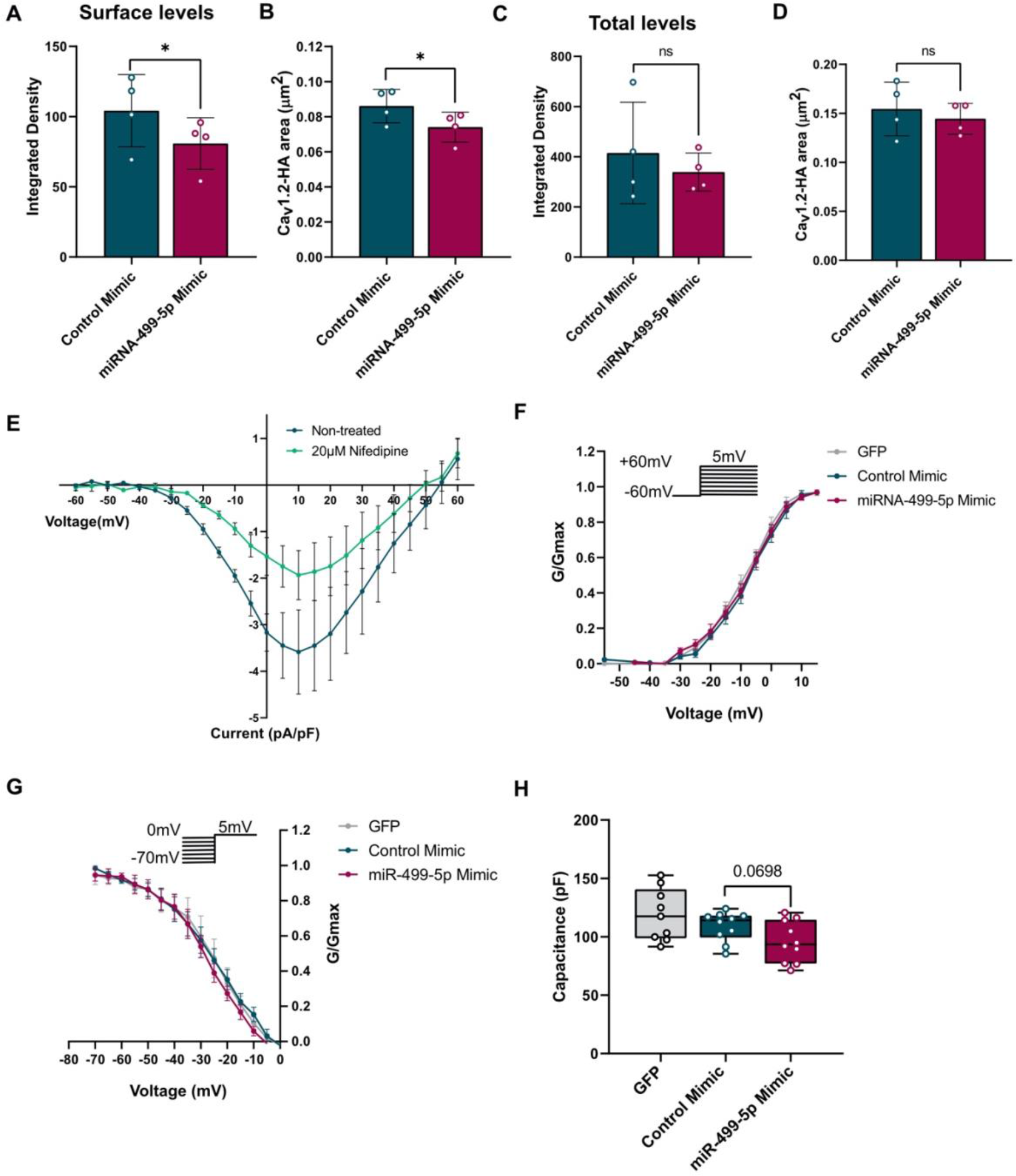
Neurons transfected with miR-499-5p mimic showed significant reductions in A) the integrated density of surface Cav1.2-HA (n=4; Paired two-sample t-Test *p=0.0385) and B) the area of surface Cav1.2-HA (n=4; Paired two-sample t-Test *p=0.0381). No significant changes were found for C) the integrated density (n=4; Paired two-sample t-Test *p=0.3213) and D) the area (n=4; Paired two-sample t-Test *p=0.3182) of total Cav1.2 channels. E) I/V curves before and after bath application of 20 μM of the L-type blocker Nifedipine (Calbiochem). F) Hippocampal neurons transfected with the miR-499-5p mimic did not show a different activation curve compared to the activation curves of cells transfected with the Control mimic or GFP alone. G) Hippocampal neurons overexpressing miR-499-5p did not show a different inactivation curve compared to the inactivation curves of control cells. H) MiR-499-5p overexpression tended to decrease capacitance.

**Supplementary Figure 5:**
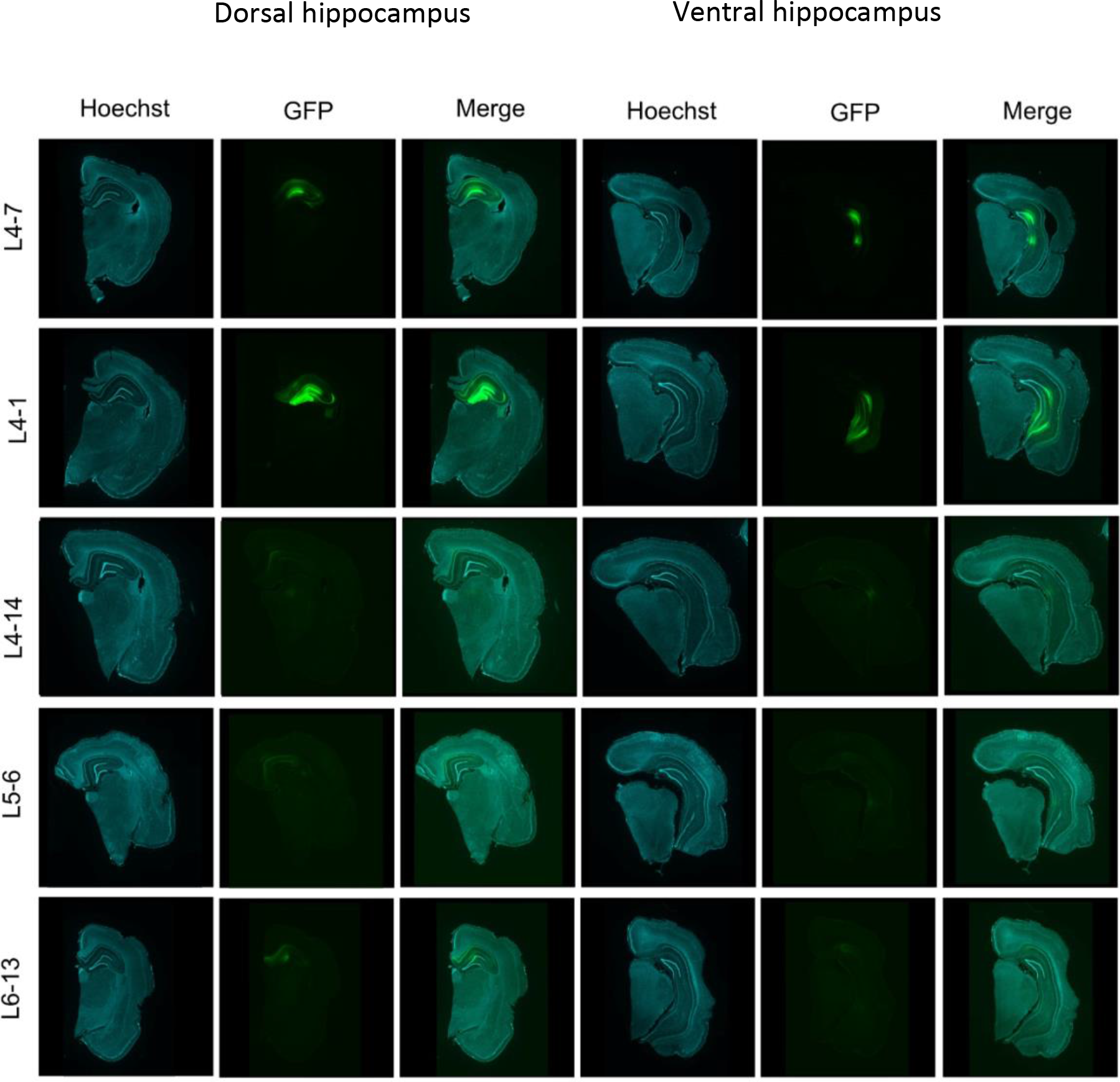
Validation of hippocampal rAAV expression by the presence of GFP fluorescence in the left hemisphere of rats expressing chimeric hairpins. Left hemispheres were dissected from rats injected bilaterally in the dorsal and ventral hippocampus with either miRNA-499 hairpin or control hairpin. Coronal brain sections of 80 μm were stained with Hoechst. Green fluorescence represents the GFP native signal of the chimeric hairpin. L4-7: left hemisphere of a rat injected with the chimeric control hairpin showing strong GFP fluorescence in both dorsal and ventral hippocampus; L4-1: left hemisphere of a rat injected with the chimeric miR-499 hairpin showing strong GFP fluorescence in both dorsal and ventral hippocampus; L4-14, L5-6 and L6-13: left hemisphere of rats injected with the chimeric miR-499 hairpin showing strong GFP fluorescence in both dorsal and ventral hippocampi.

**Supplementary Figure 6:**
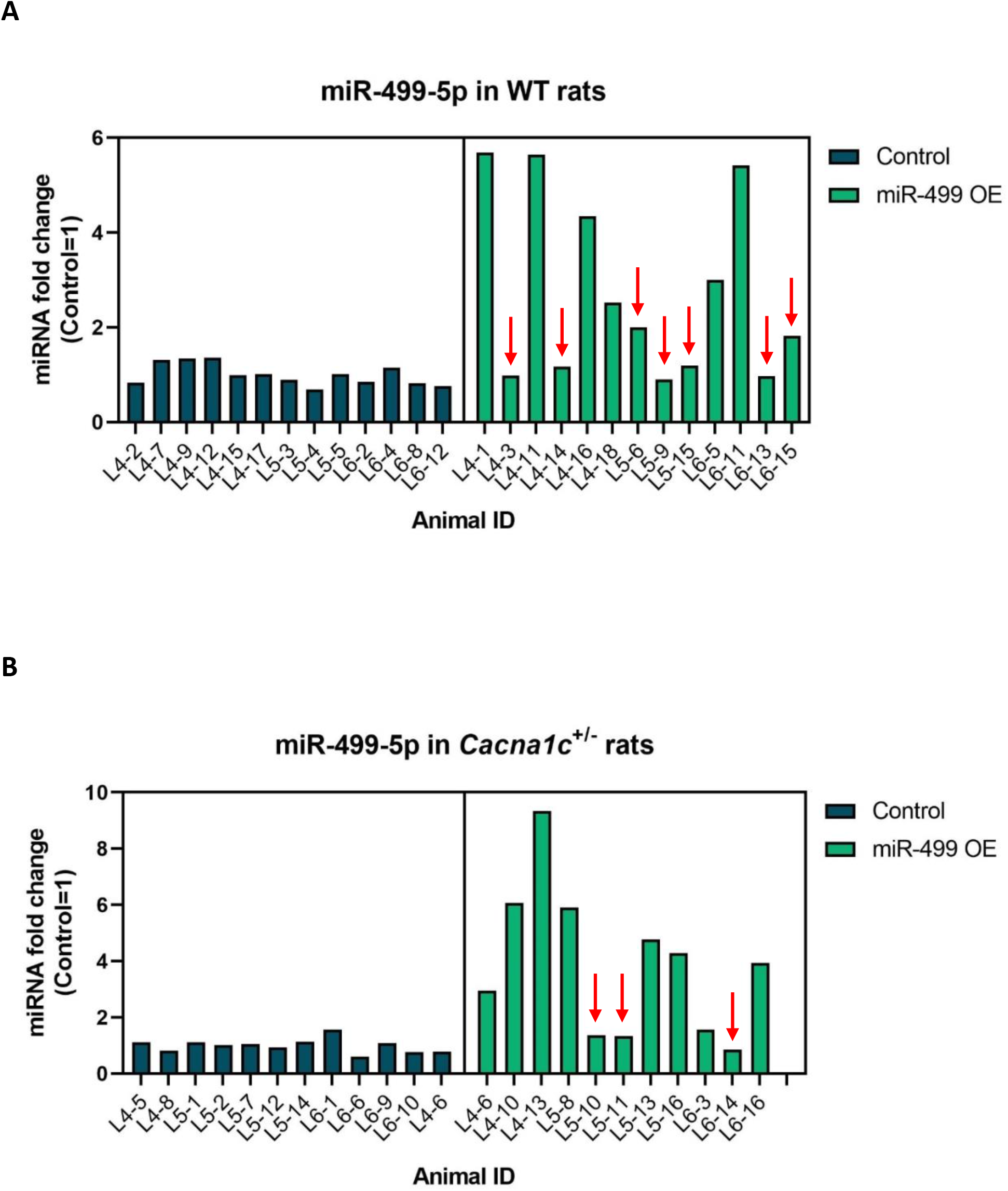
Validation of miR-499-5p overexpression in vivo using RT-qPCR. A) and B) Fold expression of miR-499-5p in WT rats relative to control sequence expression from chimeric hairpins. Rats of 1.5-2 months old were injected bilaterally in the dorsal and ventral hippocampus with 1 μL of virus per injection (either AAV-Control or AAV-miR-499). At approximately 3 months of age, rats were killed, and the brains were removed. Hippocampi dissected from the right hemisphere were freshly snap-frozen and stored at −80°C until lysis for RNA isolation and real-time qRT-PCR. Red shaded rows and arrows highlight animals that were excluded from biochemical and behavioral analysis because miR-499-5p expression was lower than the average miR-499-5p expression of animals injected with the control hairpin + 2 s.d. C) Fold expression of miR-499-5p in *Cacna1c*^+/−^ rats relative to control sequence expression from chimeric hairpins. Rats were injected as described before. Red arrows highlight animals that were excluded from behavioral analysis since miR-499-5p expression was lower than the average miR-499-5p expression of animals injected with the control hairpin + 2 s.d. For behavioral phenotyping and Western blot analysis, the following animals injected with the chimeric miR-499 hairpin were excluded due to poor miR-499 overexpression in the right hippocampus: L4-3, L4-14, L5-6, L5-9, L5-15, L6-13, and L6-15. All the animals injected with the control hairpin were included in the analysis. For Western blot, the following animals injected with the control hairpin were processed for Western blot: L4-2, L4-9, L4-12, L4-15, L4-17, L5-3, L5-4 and L5-5.

**Supplementary Figure 7:**
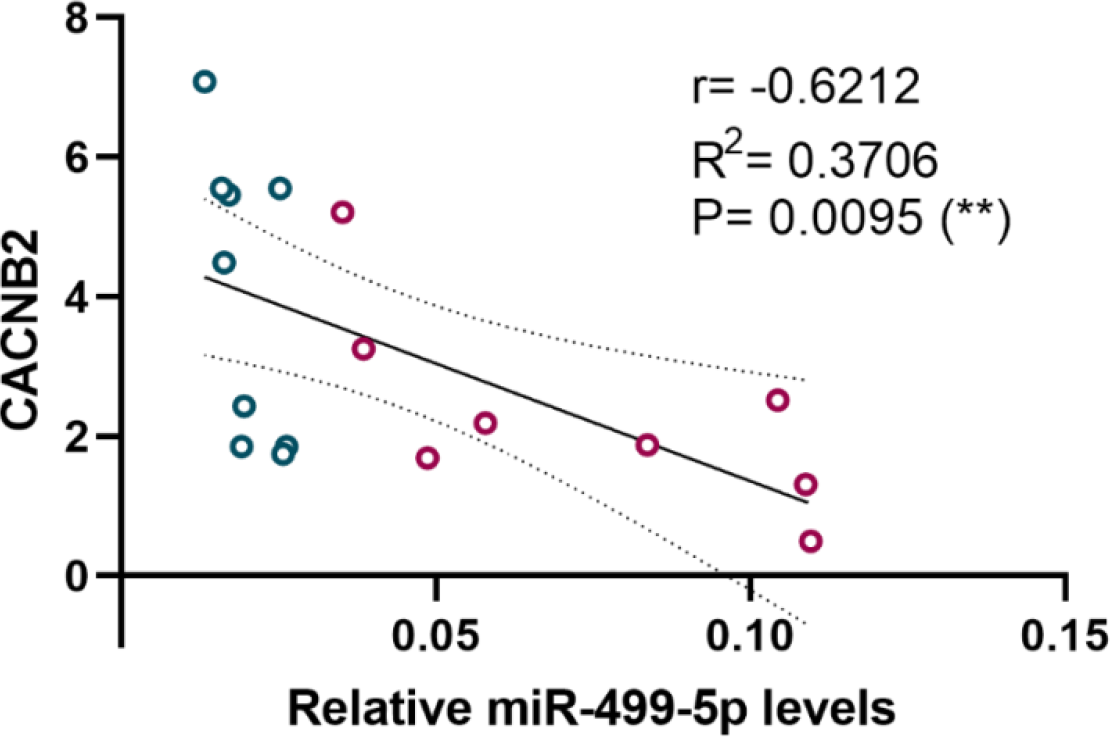
Significant positive correlation between relative miR-499-5p levels in the rat hippocampus and the protein levels of CACNB2.

